# Sex-Specific Development in the Persistence of Stress-Induced Enhancement of Fear Learning in Juvenile and Adult Rats

**DOI:** 10.64898/2026.10.04.756589

**Authors:** Samantha R. Hetherington, Andrew M. Poulos

## Abstract

It is estimated that approximately fifty percent of the population will encounter at least one traumatic event during their lifetime; however, only six percent develop Posttraumatic Stress Disorder (PTSD), a trauma-related disorder characterized by persistent alterations in affect and cognition, including an exaggerated fear response. It is well established that females (9.7%) are more than twice as likely as males (3.6%) to be diagnosed with PTSD. Nevertheless, a significant challenge remains in understanding how the severity of traumas experienced in childhood versus adulthood influences PTSD development across sexes. To address this challenge effectively, the adoption of preclinical animal models that accurately reflect these key characteristics is crucial. The Stress Enhanced Fear Learning (SEFL) paradigm in rodents encapsulates some of these complexities associated with PTSD, such as persistent fear sensitization following exposure to acute trauma. This study investigates how stress severity and developmental timing affect the long-term maintenance of SEFL in male and female rats. Juvenile and adult rats were initially subjected to no stress (control, context only), low stress (1 footshock), moderate stress (3 footshocks), or high stress (15 footshocks), followed by an extinction session either one day (recent) or sixty days (remote) later. Subsequently, to evaluate SEFL in a new context, all subjects received a single footshock and were tested the following day. Shortly after stress induction, all groups displayed SEFL. Although the effects persisted for sixty days in adult males and juvenile females, even at the lowest stress level, they were significantly reduced in juvenile males and adult females. These findings offer novel insights into sex-related differences in the persistence of SEFL and introduce new considerations regarding how the timing of developmental stressors influences future threat encoding and fear responses.

## Introduction

A single threat event rapidly engages a fear system that supports adaptive learning, allowing organisms to respond appropriately to danger (Bolles, 1970; Fanselow, 1986). Under typical conditions, fear responses are proportional to the magnitude and predictability of the threat encountered (Blanchard & Blanchard, 1969; Fanselow, 1986). However, prior exposure to uncontrollable or unpredictable stress can dysregulate these systems, producing exaggerated and persistent fear responses that generalize beyond the original traumatic experience (Nishimura et al., 2022; Deslauriers et al., 2018; Rau, & Fanselow, 2009; Poulos et al 2014). In humans, maladaptive fear regulation is considered a defining feature of anxiety- and trauma-related disorders, including Post-Traumatic Stress Disorder (PTSD), which disproportionately affects females relative to males (Kessler et al., 2005; Kilpatrick et al., 2013).

PTSD develops following exposure to traumatic events and is characterized by enduring alterations in cognition, affect, arousal, and behavior, including persistent hypervigilance and enhanced fear responsivity that endure beyond the immediate post-trauma period (American Psychiatric Association, 2013; Kessler et al., 2005). Epidemiological studies estimate that approximately 6.8% of adults in the United States will meet diagnostic criteria for PTSD during their lifetime, with prevalence rates more than twice as high in females compared with males (Kessler et al., 2005; Harvard Medical School, 2007; Kilpatrick et al., 2013). Similar sex differences emerge during adolescence, where trauma-exposed females exhibit substantially greater rates of PTSD diagnosis than males (Merikangas et al., 2010; McLaughlin et al., 2013; Cisler & Herringa, 2021). Although prevalence estimates during early childhood remain less clearly defined, accumulating evidence suggests that vulnerability to trauma-related psychopathology varies across development (Elklit et al., 2019). Together, these findings underscore the importance of identifying developmental and sex-specific mechanisms that contribute to trauma susceptibility and resilience (Kilpatrick et al., 2013; McLaughlin et al., 2013). Preclinical models have been instrumental in elucidating the neurobiological substrates of trauma-related disorders. Several rodent paradigms, including single prolonged stress, chronic variable stress, and social defeat procedures, reproduce behavioral and physiological features relevant to PTSD (Bowers et al., 2019; Flandreau & Toth, 2017; Richter-Levin, Stork, & Schmidt, 2019). However, many of these models provide limited experimental control over trauma severity and the long-term persistence of sensitized fear responses (Perusini et al., 2016). The stress-enhanced fear learning (SEFL) model addresses these limitations by employing an acute trauma session consisting of unpredictable footshocks that subsequently sensitize fear learning to a novel environment (Nishimura et al., 2022; Rajbhandari, Gonzalez, & Fanselow, 2018; Rau, & Fanselow, 2009). SEFL produces robust and persistent fear sensitization that can endure for months following trauma exposure in adult male rodents (Rau, & Fanselow, 2009). Importantly, the model permits systematic manipulation of trauma magnitude by varying shock number and enables assessment of fear persistence (Poulos et al., 2015; Rau et al., 2005) across distinct developmental time points, making it particularly well suited for examining factors that modulate long-term trauma outcomes (Poulos et al., 2015; Rau & Fanselow, 2009).

Despite the growing use of the SEFL paradigm, most studies have focused primarily on adult male subjects (Deslauriers et al., 2018; Nishimura et al., 2022; Richter-Levin, Stork, & Schmidt, 2019). Although several reports indicate that males and females exhibit comparable levels of SEFL immediately following trauma exposure (Hassien et al., 2020; Poulos et al., 2015), emerging evidence suggests that downstream behavioral consequences may diverge by sex (Gonzalez et al., 2021). For example, trauma exposure has been associated with increased alcohol consumption in males but elevated baseline anxiety-like behavior in females following SEFL induction (Gonzalez et al., 2021). Whether trauma severity differentially influences the long-term persistence of SEFL in females relative to males remains poorly understood. Moreover, developmental investigations of SEFL are comparatively limited. Stress exposure during infancy has been shown to produce sensitized fear responses that persist into juvenile and adult periods (Poulos et al., 2014; Quinn, Skipper, Claflin, 2014), and recent findings suggest that late-infancy trauma induces persistent SEFL in females but not males, raising the possibility of sex-specific sensitive periods in trauma susceptibility (Minshall et al., 2024). These observations align with broader developmental literature demonstrating that normative fear learning and regulation differ across age and sex, with males generally exhibiting increasing contextual fear across maturation, whereas females display the opposite trajectory (Colon et al., 2018; Maren et al., 1994).

Collectively, these findings highlight critical gaps in understanding how sex, developmental stage, and trauma severity interact to shape the persistence of trauma-related fear sensitization. The present study examined whether varying levels of stress induction modulate the magnitude and persistence of SEFL in juvenile (P24) and adult (P90) male and female rats. Animals were exposed to no-shock, low-intensity (1-shock), moderate-intensity (3-shock), or high-intensity (15-shock) trauma conditions, followed by extinction procedures and assessment of fear sensitization in a novel context at either recent (1 day) or remote (60 days) intervals. We hypothesized that both adult males and females would exhibit SEFL at recent time points, but that sex differences would emerge during remote testing. In juvenile subjects, we predicted that both sexes would develop persistent SEFL, although trauma severity was expected to differentially influence the stability and magnitude of sensitized fear responses across development.

## Methods

### Subjects

Adult (P90) and juvenile (P24) male and female Long Evans rats bred in the Life Sciences Animal Facility at the University at Albany were used for the following experiments (breeders originally purchased from Taconic Biosciences, Inc.). All rats were housed in a 12-hour light-dark cycle with lights on from 8:00 AM to 8:00 PM in a temperature (70° +/-2° F) controlled vivarium. Rats were housed in Ancare clear polycarbonate cages (Ancare, Bellmore, NY; 10 ½" x 19” x 8”) that included a stainless-steel wire lid and Micro-Tilter Top™ with wood shavings for bedding and *ad libitum* access to water and rat chow (5P76/Isopro RMH 3000) from Lab Diet (Adred Hills, MN).

The day of birth was designated as postnatal day (P) zero. At P21, all subjects were weaned from dams’ home cages and housed in age and sex-matched cages of 2 to 4 littermates. After three days of handling, testing began at P24 (juvenile) or P90 (adult). All animals were tested between 12:00 PM and 6:00 PM. All behavioral experiments were conducted by a cis-female investigator.

### Apparatus

All behavioral experiments were conducted in light and sound-controlled Med Associates fear conditioning chambers (MED-VFC2-USB-R w/VFC-008-LP. Saint Albans, VT). All stimuli were programmed and controlled with computers (Seneca, PRO437526, Microsoft Windows 7 Professional) that ran Med Associates VideoFreeze Software (SOF-843). A near-infrared camera (VID-CAM-MONO-4) affixed to the chamber door recorded rodent behavior during each session. Each chamber housed a conditioning box with clear plexiglass front, back, ceiling, and aluminum sidewalls. Conditioning boxes were configured into two conditioning contexts (A, B) that differed based on visual, olfactory, tactile, and auditory features.

Context A consisted of an evenly spaced grid floor made from stainless steel rods measuring either 1/8 (VFC-005A for P24 subjects) or 3/16 (VFC-005 for P90 subjects) in diameter. A 50% simple green solution was sprayed into drop pans as a scent for Context A. Background noise was measured at 63 dB. Context B consisted of a vertically staggered grid floor with rods measuring either 1/8 (VFC-005A-S for P24 subjects) or 3/16 (VFC-005-S for P90 subjects) in diameter and an A-Frame-shaped ceiling and side walls (VFC-008-IRT). While there was no lighting in the chamber, a standing lamp with a red incandescent bulb was used to illuminate the room so that the investigator could maneuver. A 1% acetic acid solution was sprayed into drop pans as a scent for context B. Background noise was measured at 49 dB. Footshocks (1 mA, 2 sec) in both contexts were delivered through grid floors connected to a scrambled shock generator (ENV-414S). Both contexts, including the grid floors, were cleaned with Clidox-S at the start and end of each session.

### Measure of Freezing

Freezing was defined as the absence of movement, except for that required for respiration (Fanselow & Bolles, 1979). VideoFreeze Software (SOF-843) measures motion by determining the change in video pixel composition over time. The system measures motion on a scale from 0 to 76,800 au, with a recommended activity threshold of 10 to 20 au. The motion threshold for this experiment was set at 18 au.

### Handling

Habituation to handling has been associated with reduced stress to the handler, increased animal welfare, and more valid research (Beane et al., 2002; Madruga et al., 2006; Rajbhandari, Gonzalez, & Fanselow, 2018). Thus, all subjects were handled for four minutes per day for three days before stress induction in an empty room.

### SEFL Procedure

The SEFL procedure consisted of a four-day procedure, with two days in context A and two days in novel context B (Figure 1). The first day was referred to as stress induction, followed the next day by a 20-minute test of stress-induction memory and extinction in context A. The last two days consist of a single-trial fear conditioning and testing protocol in context B.

**Figure 1:**
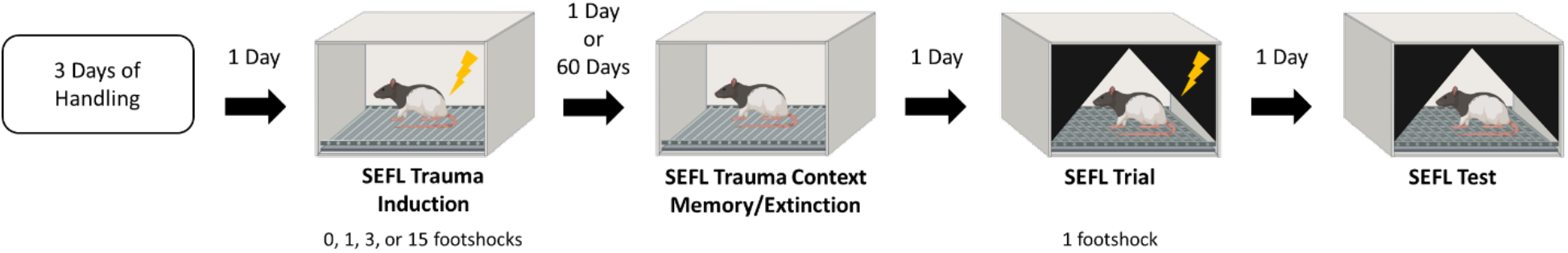
Procedural design. All subjects received three days of handling before behavioral testing, either at P21 for juveniles or P87 for adults. Then, on P24 and P90, subjects began behavioral testing with stress trauma induction of 0, 1, 3, or 15 footshocks in context A. Extinction and the context memory test took place either the next day (recent interval) or 60 days later (remote interval). The next day, subjects were placed in context B where they received a single footshock for the SEFL trial. The last day of testing consisted of a SEFL test. The following graphs use the following symbols to depict significance. An asterisk and line indicate a significant difference between the groups. A star indicates a significant difference from the no stress/control group. A triangle indicates a significant difference from the low stress group. A square indicates a significant difference from the moderate stress group. A diamond indicates a significant difference from the high stress group.

**Figure 2:**
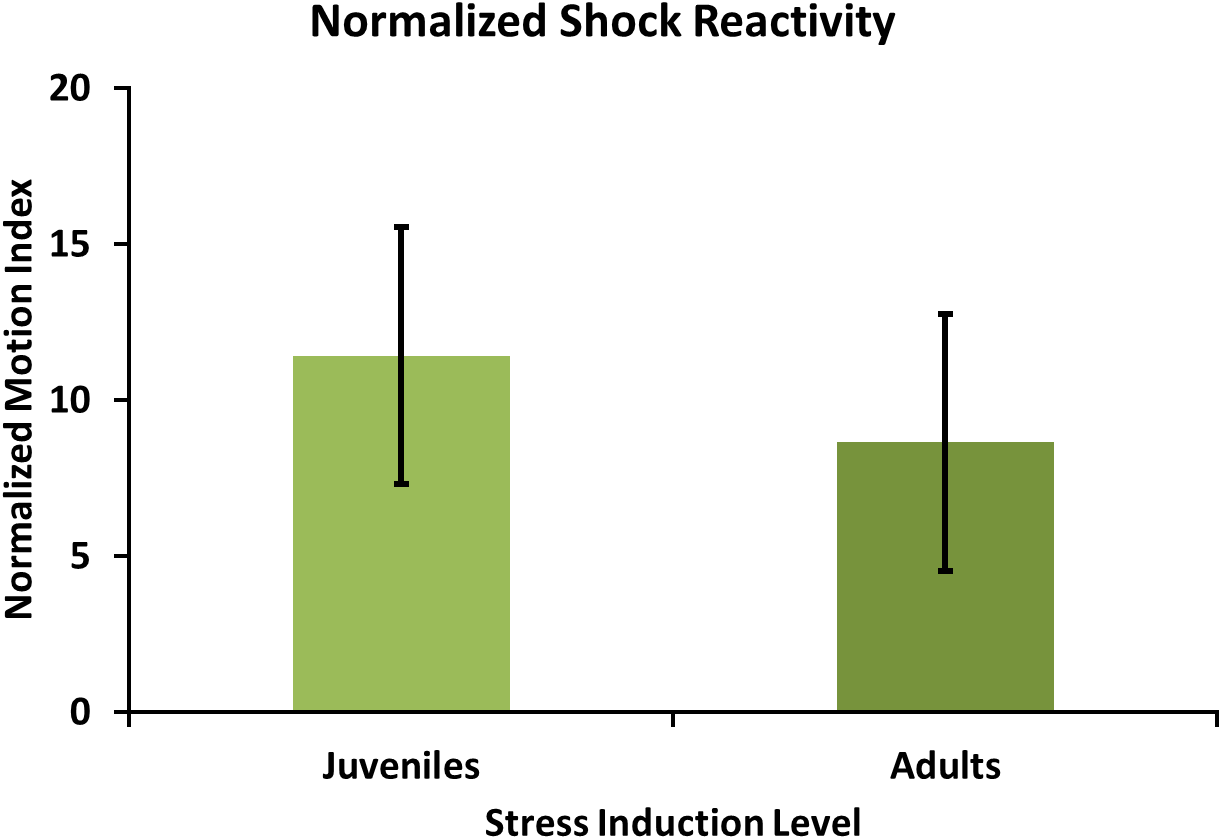
Normalized shock reactivity for juveniles and adults during stress induction in context A, as depicted by the max motion index for the 2 seconds during the first footshock, normalized to the 2 seconds prior to the first footshock. The 2×2×2×3 ANOVA that found no significant main effects or interactions for juveniles or adults.

#### Stress Trauma Induction in Context A

On day 1, subjects were transported from the vivarium to the same holding room used during handling and left to habituate to the room for 10 minutes. Subjects were then placed individually into context A conditioning boxes. Subjects underwent stress induction consisting of either 0, 1, 3, or 15 footshocks (1 mA, 2 sec), designated as no stress, low, moderate, and high stress induction, respectively (Table 1). All footshocks were delivered pseudo-randomly between three to seven minutes apart. The first three minutes and the last minute were shock-free for all group conditions. For the number of subjects per group see Table 2.

**Table 1:** Describes the number of footshocks for each stress group as well as the total duration spent in Context A. The first three minutes and the last minute were shock-free for all groups with all shocks delivered pseudo-randomly between three and seven minutes. Animals in the no stress group were divided into three groups that time-matched one of the stress groups in context A.

| Stress Group | Number of Shocks | Total Time in Context |
| --- | --- | --- |
| No Shock | 0 | 4 mins and 2 secs, 16 mins and 6 secs, or 76 mins and 30 secs |
| Low | 1 | 4 mins and 2 secs |
| Moderate | 3 | 16 mins and 6 secs |
| High | 15 | 76 mins and 30 secs |

**Table 2:** Number of animals used per group.

| Sex | Age | Interval | Stress | n |
| --- | --- | --- | --- | --- |
| Male | Adult | Recent | No | 14 |
|  |  |  | Low | 10 |
|  |  |  | Moderate | 10 |
|  |  |  | High | 8 |
|  |  | Remote | No | 14 |
|  |  |  | Low | 8 |
|  |  |  | Moderate | 8 |
|  |  |  | High | 8 |
|  | Juvenile | Recent | No | 15 |
|  |  |  | Low | 9 |
|  |  |  | Moderate | 8 |
|  |  |  | High | 8 |
|  |  | Remote | No | 13 |
|  |  |  | Low | 8 |
|  |  |  | Moderate | 9 |
|  |  |  | High | 7 |
| Female | Adult | Recent | No | 12 |
|  |  |  | Low | 10 |
|  |  |  | Moderate | 8 |
|  |  |  | High | 8 |
|  |  | Remote | No | 12 |
|  |  |  | Low | 9 |
|  |  |  | Moderate | 9 |
|  |  |  | High | 8 |
|  | Juvenile | Recent | No | 13 |
|  |  |  | Low | 8 |
|  |  |  | Moderate | 8 |
|  |  |  | High | 6 |
|  |  | Remote | No | 12 |
|  |  |  | Low | 10 |
|  |  |  | Moderate | 9 |
|  |  |  | High | 9 |

Shock reactivity will be assessed by comparing the Max motion index for the 2 seconds during the first footshock. To account for the pixel change in the software due to the size difference between adults and juveniles, the max motion index for each group will be normalized to the max motion index value for the 2 seconds prior to the first footshock. Shock conditioning for the moderate- and high-stress groups will be assessed by comparing percent freezing for the 30 seconds post each shock.

#### Stress Trauma Induction Memory Test & Extinction in Context A

Day 2 occurred either the following day (recent) or 60 days later (remote). Subjects were transported from the vivarium to the holding room and left for 10 minutes. Subjects were returned to context A for a combined 20-minute context fear test and extinction session.

The retrieval of the induction memory was assessed by percent freezing during the first 8 minutes of the session. To assess extinction rates, the percent freezing value for the first 4 and last 4 minutes of the 20-minute session will be compared.

#### SEFL Trial in Context B

Twenty-four hours after testing in context A, subjects were transported from the vivarium to a holding room with completely covered cages. They were placed in the holding room with the lights off and habituated for 10 minutes. Subjects were taken into the testing room while still covered, then placed individually into context B conditioning boxes, where they received a single footshock (1 mA, 2 sec) regardless of the initial stress induction.

To assess generalized fear in context B, the average percent freezing for the first 3 minutes before the single shock was compared.

#### SEFL Test in Context B

The following day, subjects returned to context B in the same manner as described above (SEFL Trial in Context B) for an 8-minute retrieval memory test, with no footshocks administered to any group.

Subjects will be assessed for context fear and SEFL by comparing percent freezing over the full 8 minutes of the session. To account for differences related to sex and development in context fear conditioning (Colon et al., 2018), respective control groups, a percent change score was computed, where subjects’ data were calculated as the percent change from the average of their controls.

### Statistical Analysis

Normality, equality of variance, and independence of factors we assessed for each test. A power analysis was performed for all groups *a priori* to determine suggested n values. All statistics were performed using SPSS Statistics 22 (IBM), with significance set at p < 0.05.

Groups were compared using a 2×2×2×4 ANOVA with factors for sex (male/female), age (juvenile/adult), interval (recent/remote), and stress induction (no/low/moderate/high). When applicable, a 2×2×2×3 ANOVA with factors for sex (male/female), age (juvenile/adult), interval (recent/remote), and stress induction (low/moderate/high), excluding controls. Otherwise, a two-way ANOVA was performed using between-subjects factors (stress (no, low, moderate, and high) and interval (recent and remote) for each sex and age group. If an interaction was identified, a one-way ANOVA was performed for each interval. When applicable, a repeated-measures ANOVA will be applied with between-subject factors (stress (no, low, moderate, and high) and interval (recent and remote)) for each sex and age group and within-subject factors (minutes (first four and last four) or shock (shock number)). If sphericity was not met, then Greenhouse-Geisser corrections were used. *Post-hoc* multiple comparisons were completed using a Bonferroni corrected t-test or paired t-test.

Four animals were excluded from the statistical analysis due to equipment failure.

## Results

### Normalized Shock Reactivity during Stress Induction in Context A

To evaluate potential variations in sensitivity to footshock during stress induction between male and female juvenile and adult rats, we analyzed baseline reactivity to the initial footshock for each group. To control for differences in size and initial activity levels among juveniles as well as male and female adults, individual shock-elicited motion was normalized relative to each animal’s pre-shock maximum motion index. A 2 (sex) x 2 (age) x 2 (interval) x 3 (stress severity) ANOVA for shock reactivity indicated no significant interactions or main effects (see Table 3).

**Table 3:** Statistical outcomes for normalized shock reactivity. A 2×2×2×3 ANOVA found no significant differences when accounting for size differences between juveniles and adults.

| Normalized Shock Reactivity |  |  |  |
| --- | --- | --- | --- |
| Source | F | Sig. Value | Power |
| Age | 0.196 | 0.658 | 0.073 |
| Sex | 1.464 | 0.228 | 0.225 |
| Interval | 0.045 | 0.833 | 0.055 |
| Stress Induction Severity | 0.354 | 0.703 | 0.106 |
| Age * Sex | 0.174 | 0.677 | 0.070 |
| Age * Interval | 2.019 | 0.157 | 0.293 |
| Age * Stress Induction Severity | 1.041 | 0.355 | 0.230 |
| Sex * Interval | 0.032 | 0.859 | 0.054 |
| Sex * Stress Induction Severity | 0.787 | 0.457 | 0.183 |
| Interval * Stress Induction Severity | 1.143 | 0.321 | 0.249 |
| Age * Sex * Interval | 1.104 | 0.295 | 0.181 |
| Age * Sex * Stress Induction Severity | 0.982 | 0.377 | 0.219 |
| Age * Interval * Stress Induction Severity | 0.540 | 0.584 | 0.138 |
| Sex * Interval * Stress Induction Severity | 1.782 | 0.171 | 0.370 |
| Age * Sex * Interval * Stress Induction Severity | 0.533 | 0.588 | 0.137 |

### Post-Shock Freezing during Stress Induction in Context A

To assess initial fear to stress induction procedures, we analyzed post-shock freezing separately for each stress induction level (Figure 3). Individual ANOVAs for post-shock freezing in low, moderate, and high stress induction conditions found no main effects or interactions of sex, age, or interval interval (Table 4). In the moderate (F (2, 60) = 7.751, p < 0.001) and high (F (14, 41) = 2.794, p < 0.001) stress induction conditions, between-subjects (repeated measures) analysis showed that post-shock freezing increased with repeated footshock.

**Figure 3:**
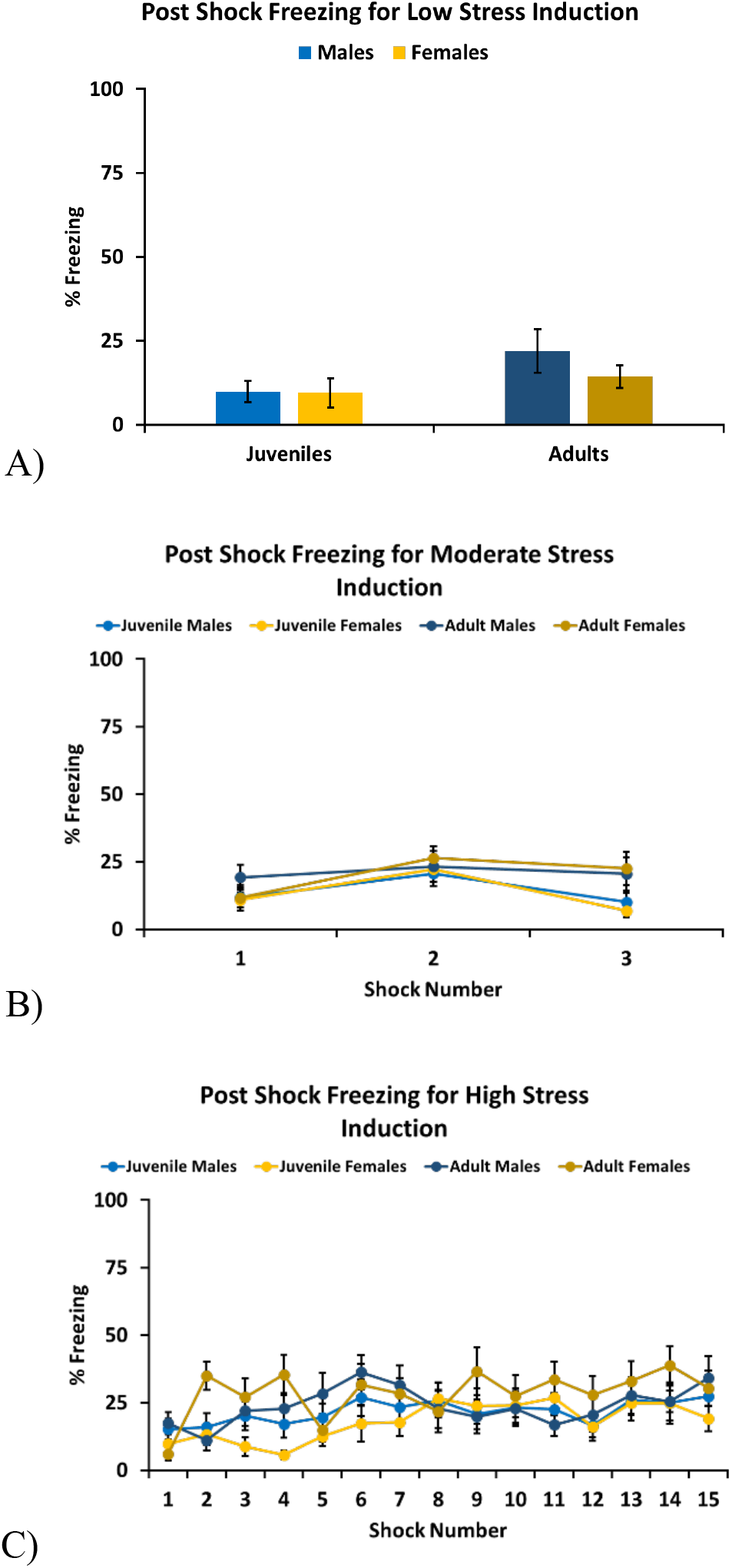
Conditioning of the group to the shocks was assessed by examining the percent freezing levels for the 30 seconds post-shock. Conditioning was only impacted by stress severity, as indicated by a 2×2×2 ANOVA for the **A)** low stress group, which found no significant differences between groups and 2×2×2 repeated ANOVAs for the **B)** moderate (F (2, 60) = 7.751, p < 0.001) and **C)** high (F (14, 41) = 2.794, p < 0.001) stress groups, which found only a main effect for shock number.

**Table 4:** Statistical outcomes for shock conditioning. A 2×2×2 ANOVA for the low stress group found no significant differences between groups and separate repeated 2×2×2 repeated ANOVAs for the moderate and high stress groups found only a main effect for shock number. For moderate stress sphericity was assumed due to a non-significant Mauchley’s test (W = 0.953, p = 0.236), but for high stress Mauchley’s test was significant (W = 0.034, p < 0.001) and thus Greenhouse-Geisser corrections were made.

| Shock Conditioning, Low Stress |  |  |  |
| --- | --- | --- | --- |
| Source | F | Sig. Value | Power |
| Age | 3.182 | 0.079 | 0.420 |
| Sex | 0.799 | 0.375 | 0.142 |
| Interval | 0.279 | 0.599 | 0.081 |
| Age * Sex | 0.726 | 0.397 | 0.134 |
| Age * Interval | 0.026 | 0.873 | 0.053 |
| Sex * Interval | 3.518 | 0.065 | 0.455 |
| Age * Sex * Interval | 1.380 | 0.244 | 0.212 |

| Shock Conditioning, Moderate Stress |  |  |  |
| --- | --- | --- | --- |
| Source | F | Sig. Value | Power |
| Shock Number | 7.751 | < 0.001 | 0.946 |
| Shock Number * Age | 2.310 | 0.104 | 0.461 |
| Shock Number * Sex | 0.758 | 0.471 | 0.176 |
| Shock Number * Interval | 0.115 | 0.891 | 0.067 |
| Shock Number * Age * Sex | 0.525 | 0.593 | 0.135 |
| Shock Number * Age * Interval | 1.539 | 0.219 | 0.322 |
| Shock Number * Sex * Interval | 1.715 | 0.184 | 0.355 |
| Shock Number * Age * Sex * Interval | 1.582 | 0.210 | 0.330 |

**Table 4:** Statistical outcomes for shock conditioning.
| Shock Conditioning, High Stress |  |  |  |
| --- | --- | --- | --- |
| Source | F | Sig. Value | Power |
| Shock Number | 2.794 | 0.002 | 0.971 |
| Shock Number * Age | 1.062 | 0.390 | 0.561 |
| Shock Number * Sex | 1.557 | 0.117 | 0.766 |
| Shock Number * Interval | 0.845 | 0.584 | 0.448 |
| Shock Number * Age * Sex | 0.850 | 0.580 | 0.451 |
| Shock Number * Age * Interval | 0.663 | 0.757 | 0.349 |
| Shock Number * Sex * Interval | 1.051 | 0.399 | 0.555 |
| Shock Number * Age * Sex * Interval | 0.807 | 0.620 | 0.428 |

### Retrieval of Stress Induction Memory in Context A

To evaluate the long-term retention of the contextual fear memory of the stress induction context, a full factorial 2 (sex) x 2 (age) x 2 (interval) x 4 (stress severity) ANOVA revealed a four-way interaction (F (3, 276) = 2.638, p = 0.05) (Table 5). To further evaluate the impact of stress severity on retention interval, separate ANOVAs for adult male, adult female, juvenile male, and juvenile female rats were conducted.

**Table 5:** Full factorial statistical outcomes for retrieval of stress induction memory. A 2×2×2×4 ANOVA found an interaction between age, sex, interval, and stress induction severity. This was followed by 2×4 ANOVAs investigating interval and stress induction severity for adult males, adult females, juvenile males, and juvenile females.

| Retrieval of Stress Induction Memory |  |  |  |
| --- | --- | --- | --- |
| Source | F | Sig. Value | Power |
| Age | 36.243 | < 0.001 | 1.000 |
| Sex | 1.446 | 0.230 | 0.224 |
| Interval | 39.243 | < 0.001 | 1.000 |
| Stress Induction Severity | 314.247 | < 0.001 | 1.000 |
| Age * Sex | 5.328 | 0.022 | 0.633 |
| Age * Interval | 15.927 | < 0.001 | 0.978 |
| Age * Stress Induction Severity | 7.027 | < 0.001 | 0.980 |
| Sex * Interval | 0.039 | 0.844 | 0.054 |
| Sex * Stress Induction Severity | 0.848 | 0.469 | 0.234 |
| Interval * Stress Induction Severity | 6.212 | < 0.001 | 0.963 |
| Age * Sex * Interval | .305 | 0.581 | 0.085 |
| Age * Sex * Stress Induction Severity | 1.273 | 0.284 | 0.339 |
| Age * Interval * Stress Induction Severity | 4.749 | 0.003 | 0.897 |
| Sex * Interval * Stress Induction Severity | 1.084 | 0.356 | 0.292 |
| Age * Sex * Interval * Stress Induction Severity | 2.631 | 0.050 | 0.641 |

Adult male rats showed a robust, enduring contextual fear memory that persisted across recent and remote interval periods (Figure 4A). This was supported by a 2 (interval) x 4 (stress severity) ANOVA that found a main effect of stress induction severity (F (3, 72) = 126.621, p < 0.001), but did not find an interaction (F (3, 72) = 1.124, p > 0.05 nor a main effect for interval (F (1, 72) = 2.107, p = 0.151) (Table 6.) *Post hoc* comparisons confirmed contextual fear memory in all stress induction groups, with the high (p < 0.001), moderate (p < 0.001), and low (p < 0.001) stress groups showing significantly greater freezing than the non-stressed group. Furthermore, stress induction intensity was directly proportional to the degree of contextual fear memory. This finding was supported by the observation that the high-stress group exhibited significantly greater fear memory than the moderate (p < 0.001) and low (p < 0.001) stress groups, which also differed significantly from each other (p < 0.001).

**Figure 4:**
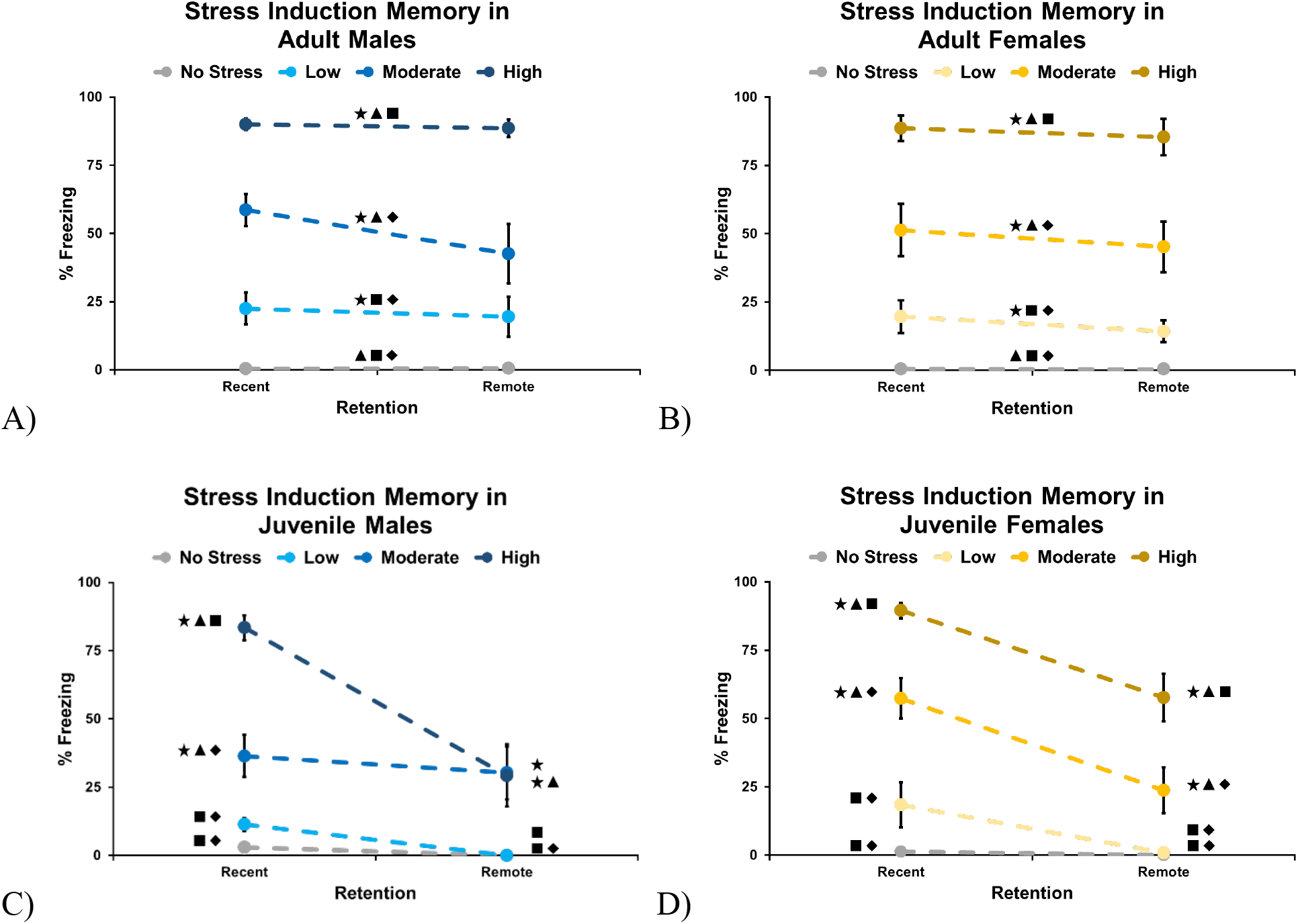
Retrieval of stress induction memory. **A)** Adult males show a robust and continued contextual fear memory that was graded by stress severity but not impacted by interval. **B)** Adult females also demonstrated a robust contextual fear memory that was impacted by stress severity but not interval. **C)** Juvenile males demonstrated a graded robust context fear memory when tested at a recent interval period but that did not persist as seen at the remote interval test period. **D)** Juvenile females demonstrated a graded robust context fear memory when tested at a recent interval period but that did not persist as seen at the remote interval test period.

**Table 6:**
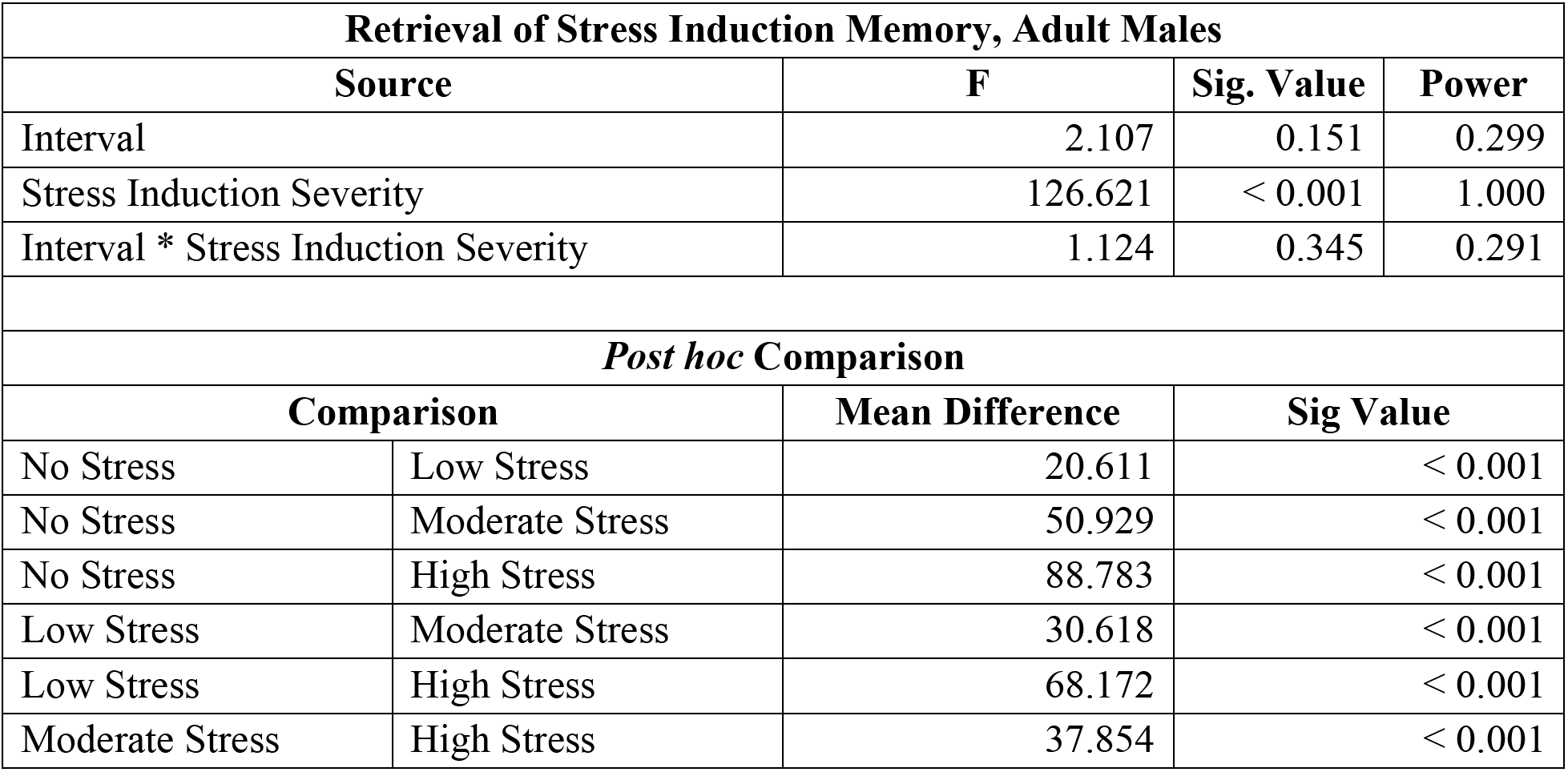
Statistical outcomes for retrieval of stress induction, adult males. Only a main effect of stress induction severity was found with all groups significantly differing from one another.

Adult female rats also demonstrated a robust contextual fear memory that persisted and did not differ between the recent and remote interval periods (Figure 4B). A 2 (interval) x 4 (stress severity) ANOVA supported this with a main effect of stress induction severity (F (1, 68) = 93.820, p < 0.001). No interaction between interval and stress induction severity (F (3, 68) = 0.138, p > 0.05) was found, nor was there a main effect of interval (F (1, 68) = 0.892, p > 0.05)(Table 7). A *post hoc* comparison between stress groups confirmed the development of a contextual fear memory in all stress induction groups, with the high (p < 0.001), moderate (p < 0.001), and low (p < 0.05) stress groups exhibiting significantly greater freezing than the no-stress group. As in adult male rats, the high-stress group was significantly greater than the moderate (p < 0.001) and low (p < 0.001) stress groups, which also differed significantly from each other (p < 0.001).

**Table 7:**
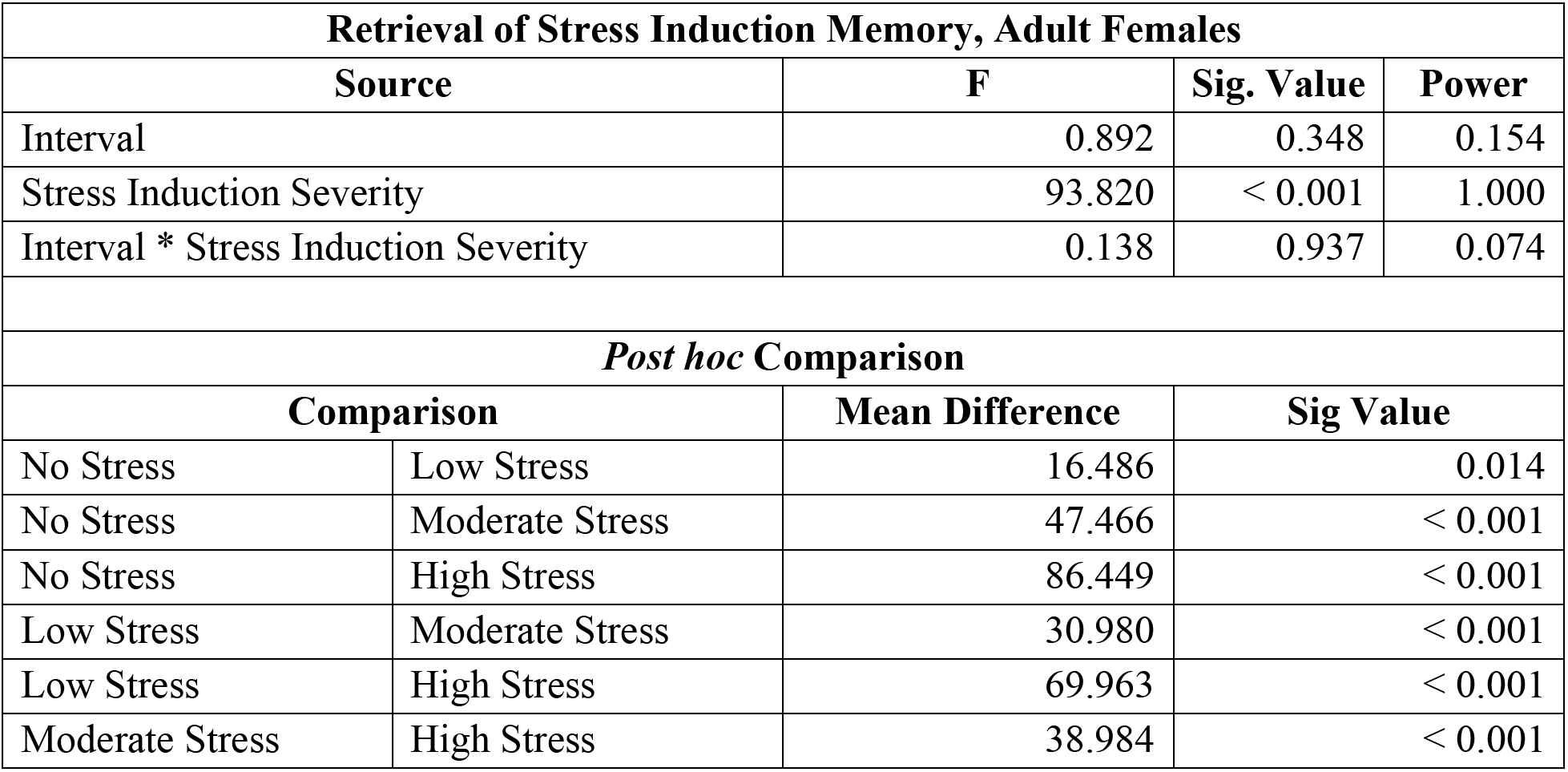
Statistical outcomes for retrieval of stress induction, adult females. Only a main effect of stress induction severity was found with all groups significantly differing from one another.

In juvenile male rats, context fear memory was robust at the recent interval period but not at the remote interval test period (Figure 4C). A 2 (interval) x 4 (stress severity) ANOVA supported this, showing an interaction between interval and stress induction severity (F (3, 69) = 8.289, p < 0.001). One-way ANOVAs followed, showing a main effect of stress severity at recent (F (3, 36) = 86.233, p < 0.001) and remote (F (3, 33) = 6.309, p > 0.01) interval (Table 8). A *post hoc* comparison between stress groups at recent interval confirmed that stress induction produced a contextual fear memory for the high (p < 0.001) and moderate (p < 0.001) stress groups, as they demonstrated significantly higher freezing than the no stress group; however, no significant difference was found between the low (p > 0.05) and no stress groups. Overall, the moderate stress group demonstrated significantly higher freezing than the low stress group (p < 0.001) and significantly lower freezing than the high stress group (p < 0.001). A *post hoc* comparison between stress groups at remote interval confirmed that stress induction produced a contextual fear memory that persisted for the high (p < 0.05) and moderate (p < 0.01) stress groups as they demonstrated significantly higher freezing than the no stress group, however no significant difference was found between the low (p > .05) and no stress groups. This effect did not appear to be graded, as there was no significant difference between the high- and moderate-stress groups (p > .05).

**Table 8:** Statistical outcomes for retrieval of stress induction, juvenile males. A 2×4 ANOVA found an interaction between interval and stress induction severity. This was followed by a one-way ANOVA for each interval period which found a main effect of stress induction at both interval periods. *Post hoc* tests demonstrate a graded effect based on stress severity for the recent but not the remote interval.

| Retrieval of Stress Induction Memory, Juvenile Males |  |  |  |
| --- | --- | --- | --- |
| Source | F | Sig. Value | Power |
| Interval | 20.657 | < 0.001 | 0.994 |
| Stress Induction Severity | 41.301 | < 0.001 | 1.000 |
| Interval * Stress Induction Severity | 8.289 | < 0.001 | 0.990 |

| Retrieval of Stress Induction Memory, Juvenile Males @ Recent Interval |  |  |  |
| --- | --- | --- | --- |
| Source | F | Sig. Value | Power |
| Stress Induction Severity | 86.233 | < 0.001 | 1.000 |
| *Post hoc Comparison* |  |  |  |
| Comparison |  | Mean Difference | Sig Value |
| No Stress | Low Stress | 8.281 | 0.656 |
| No Stress | Moderate Stress | 33.423 | < 0.001 |
| No Stress | High Stress | 80.400 | < 0.001 |
| Low Stress | Moderate Stress | 25.142 | < 0.001 |
| Low Stress | High Stress | 72.120 | < 0.001 |
| Moderate Stress | High Stress | 46.977 | < 0.001 |

**Table 8:** Statistical outcomes for retrieval of stress induction, juvenile males.
| Retrieval of Stress Induction Memory, Juvenile Males @ Remote Interval |  |  |  |
| --- | --- | --- | --- |
| Source | F | Sig. Value | Power |
| Stress Induction Severity | 6.309 | 0.002 | 0.945 |
| *Post hoc Comparison* |  |  |  |
| Comparison |  | Mean Difference | Sig Value |
| No Stress | Low Stress | 0.016 | 1.000 |
| No Stress | Moderate Stress | 31.812 | 0.009 |
| No Stress | High Stress | 29.240 | 0.037 |
| Low Stress | Moderate Stress | 31.796 | 0.025 |
| Low Stress | High Stress | 29.224 | 0.073 |
| Moderate Stress | High Stress | 2.572 | 1.000 |

In juvenile females, rats showed robust context fear memory at the recent interval period, but it did not persist at the remote interval test period (Figure 4D). A 2 (interval) x 4 (stress severity) ANOVA supported this, identifying an interaction between interval and stress severity (F (3, 67) = 4.381, p < 0.01). Separate one-way ANOVAs then found a main effect of stress induction at recent (F (3, 31) = 56.072, p < 0.001) and remote (F (3, 36) = 24.034, p < 0.001) interval (Table 9). A post hoc comparison between stress groups at recent interval confirmed that stress induction produced a contextual fear memory in the high (p < 0.001) and moderate (p < 0.001) stress groups, which showed significantly higher freezing than the no-stress group. However, the low-stress and no-stress groups did not differ significantly (p > 0.05). The high-stress group showed significantly greater freezing than the moderate-stress group (p < 0.01). Comparisons between stress groups at remote interval confirmed that stress induction produced a contextual fear memory that persisted for the high (p < 0.001) and moderate (p < 0.05) stress groups, as they demonstrated significantly higher freezing than the no-stress group; however, no significant difference was found between the low (p > 0.05) and no-stress groups. At the remote interval, the high-stress group again demonstrated significantly greater freezing than the moderate-stress group (p < 0.01).

**Table 9:**
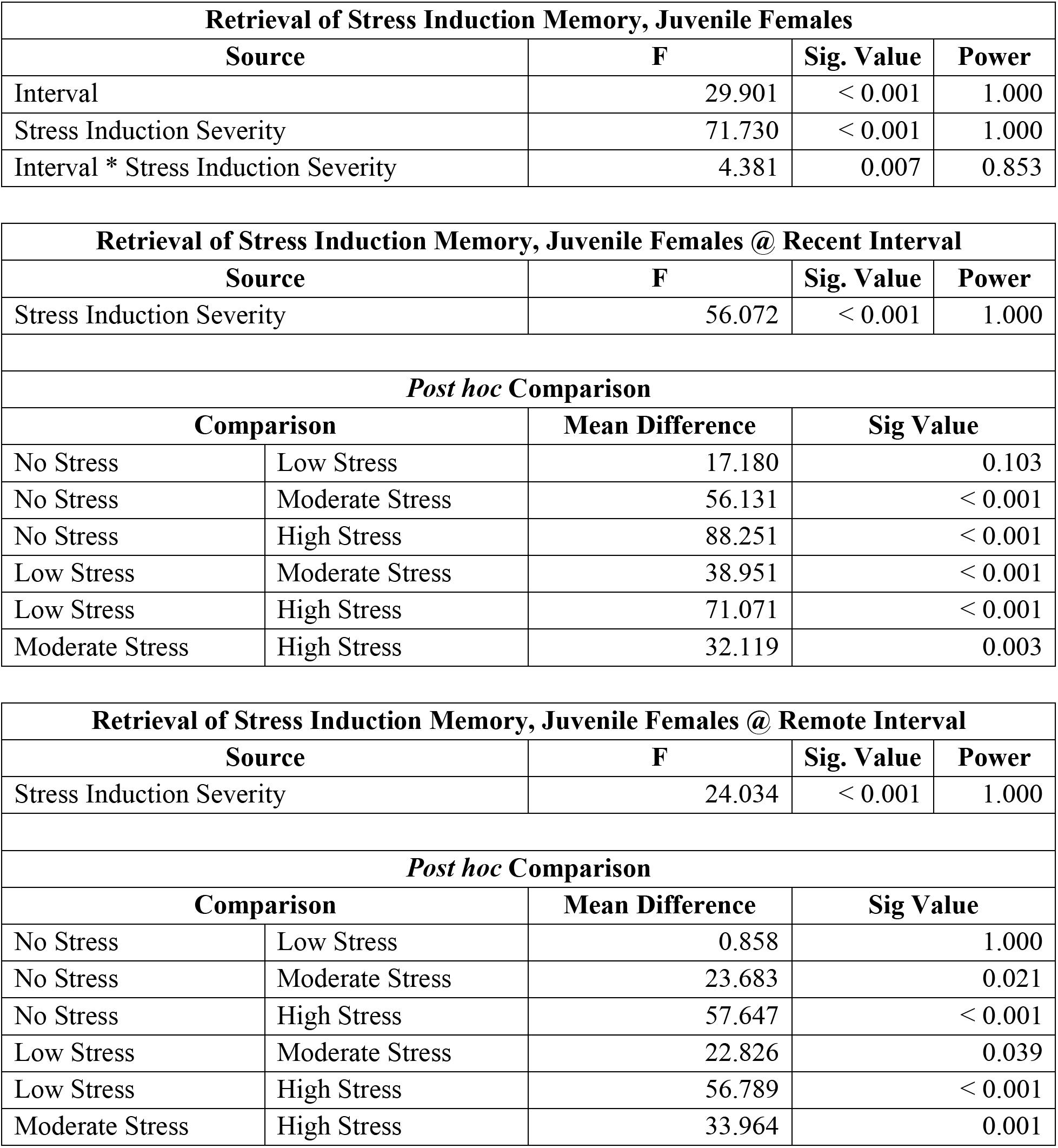
Statistical outcomes for retrieval of stress induction, juvenile females. A 2×4 ANOVA found an interaction between interval and stress induction severity. This was followed by a one-way ANOVA for each interval period which found a main effect of stress induction at both interval periods. *Post hoc* tests demonstrate a graded effect based on stress severity for the recent but not the remote interval.

### Extinction of Stress Induction Memory in Context A

Extinction of contextual fear to the stress-induction context was assessed by comparing freezing during the first four and last four minutes of the 20-minute extinction session. A repeated-measures ANOVA was performed on the within-subjects variable of time (first four and last four minutes) and between-subject variables of sex, interval, and stress severity. This repeated-measures 2 (time (first/last four minutes)) x 2 (sex) x 2 (interval) x 3 (stress severity) ANOVA revealed a main effect of time (F (1, 179) = 91.212, p < 0.001) and an interaction between time, interval, and stress severity (F (2, 179) = 7.734, p < 0.001) (Table 10). Following this, a repeated ANOVA with age and sex collapsed confirmed an interaction between time, interval, and stress induction severity (F (2, 197) = 8.644, p < 0.001) (Table 11).

**Table 10:** Full factorial statistical outcomes for extinction of stress induction memory. A 2×2×2×3 repeated ANOVA was conducted for the first/last four minutes of the extinction session with factors of age, sex, interval, and stress induction. This found an interaction between interval and stress induction on the repeated measures and between age and stress induction on the repeated measures.

| Extinction of Stress Induction Memory |  |  |  |
| --- | --- | --- | --- |
| Source | F | Sig. Value | Power |
| Minutes | 91.212 | < 0.001 | 1.000 |
| Minutes * Age | 0.004 | 0.948 | 0.050 |
| Minutes * Sex | 1.231 | 0.269 | 0.197 |
| Minutes * Interval | 7.169 | 0.008 | 0.759 |
| Minutes * Stress Induction Severity | 3.366 | 0.037 | 0.629 |
| Minutes * Age * Sex | 2.140 | 0.145 | 0.307 |
| Minutes * Age * Interval | 0.945 | 0.332 | 0.162 |
| Minutes * Age * Stress Induction Severity | 8.143 | < 0.001 | 0.957 |
| Minutes * Sex * Interval | 0.009 | 0.925 | 0.051 |
| Minutes * Sex * Stress Induction Severity | 1.237 | 0.293 | 0.267 |
| Minutes * Interval * Stress Induction Severity | 7.734 | < 0.001 | 0.947 |
| Minutes * Age * Sex * Interval | 0.491 | 0.484 | 0.107 |
| Minutes * Age * Sex * Stress Induction Severity | 1.519 | 0.222 | 0.320 |
| Minutes * Age * Interval * Stress Induction Severity | 0.365 | 0.695 | 0.108 |
| Minutes * Sex * Interval * Stress Induction Severity | 1.944 | 0.146 | 0.399 |
| Minutes * Age * Sex * Interval * Stress Induction Severity | 0.192 | 0.826 | 0.079 |

**Table 11:** Statistical outcomes for extinction of stress induction memory. A repeated ANOVA was conducted for the first/last four minutes of the extinction session with factors of interval, and stress induction with age and sex collapsed. This found an interaction between interval and stress induction on the repeated measures.

| Extinction of Stress Induction Memory |  |  |  |
| --- | --- | --- | --- |
| Source | F | Sig. Value | Power |
| Minutes | 86.703 | < 0.001 | 1.000 |
| Minutes * Interval | 7.227 | 0.008 | 0.763 |
| Minutes * Stress Induction Severity | 3.251 | 0.041 | 0.614 |
| Minutes * Interval * Stress Induction Severity | 8.644 | < 0.001 | 0.967 |

This was followed up with a repeated ANOVA looking at interval and stress induction severity at both interval periods separately which found a main effect of stress induction severity on freezing during the extinction session when tested at the recent (F (2, 98) = 4.782, p = 0.010) (Table 12) and remote (F (2, 99) = 6.600, p = 0.002) (Table 13) time points. A *post hoc* paired t-test found a significant difference in freezing levels between the first four and last four minutes for the low (p < 0.001) and moderate (p < 0.001) stress groups at the recent interval, but not for the high stress group (p = 0.310) (Figure 5a). A *post hoc* paired t-test found a significant difference in freezing levels for the first four and last four minutes for the low (p = 0.009), moderate (p < 0.001), and high (p < 0.001) stress groups at the remote interval (Figure 5b).

**Figure 5:**
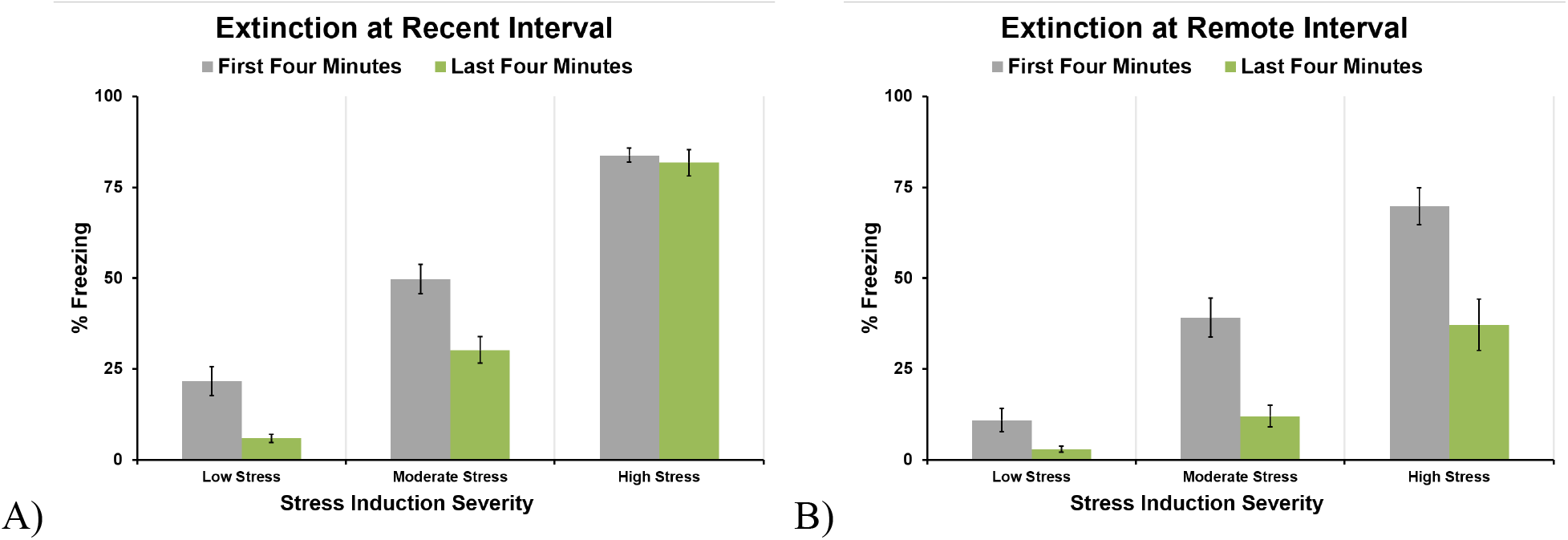
Extinction of stress induction memory as measured by freezing during the first and last 4 minutes of the 20-minute session. **A)** Freezing levels during the extinction of the stress induction memory session were found to decrease significantly for the low and moderate stress induction groups but not the high stress induction group when tested at a recent time point. **B)** Freezing levels during the extinction of the stress induction memory session were found to decrease significantly for the low, moderate, and high stress induction groups when tested at a remote time point.

**Table 12:**
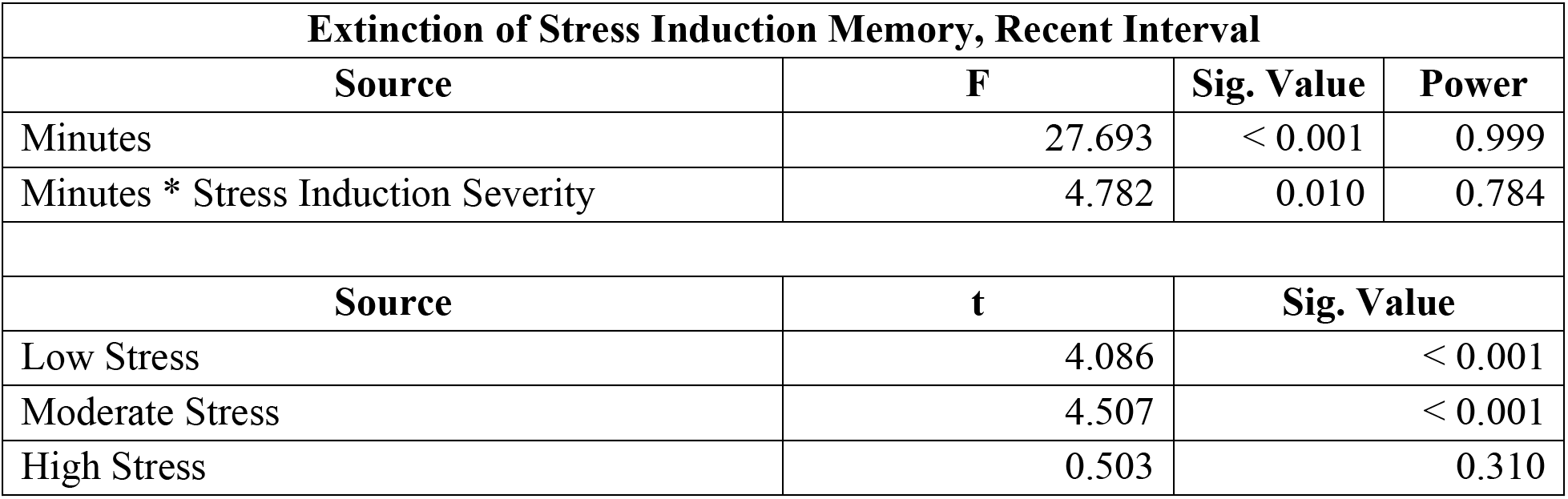
Statistical outcomes for extinction of stress induction memory. A repeated ANOVA was conducted for the first/last four minutes of the extinction session with factors of stress induction at recent interval only with age and sex collapsed.

| Extinction of Stress Induction Memory, Recent Interval |  |  |  |
| --- | --- | --- | --- |
| Source | F | Sig. Value | Power |
| Minutes | 27.693 | < 0.001 | 0.999 |
| Minutes * Stress Induction Severity | 4.782 | 0.010 | 0.784 |
| Source | t | Sig. Value |  |
| Low Stress | 4.086 | < 0.001 |  |
| Moderate Stress | 4.507 | < 0.001 |  |
| High Stress | 0.503 | 0.310 |  |

**Table 13:** Statistical outcomes for extinction of stress induction memory. A repeated ANOVA was conducted for the first/last four minutes of the extinction session with factors of stress induction at remote interval only with age and sex collapsed.

| Extinction of Stress Induction Memory, Remote Interval |  |  |  |
| --- | --- | --- | --- |
| Source | F | Sig. Value | Power |
| Minutes | 59.870 | < 0.001 | 1.000 |
| Minutes * Stress Induction Severity | 6.600 | 0.002 | 0.904 |
| Source | t | Sig. Value |  |
| Low Stress | 2.507 | 0.009 |  |
| Moderate Stress | 4.441 | < 0.001 |  |
| High Stress | 5.996 | < 0.001 |  |

### Context B Pre-Shock Stress-Induction Generalization

To assess the generalization of stress induction-related fear, freezing during the first three minutes of the SEFL trial prior to the shock in context B was examined by a full factorial 2 (sex) x 2 (age) x 2 (interval) x 4 (stress severity) ANOVA. This identified a three-way interaction between interval, age, and stress severity (F (3, 276) = 12.076, p < 0.001) and between sex, age, and stress severity (F (3, 276) = 14.644, p < 0.001) (Table 14).

**Table 14:** Full factorial statistical outcomes for the Generalized freezing in context B. A 2×2×2×4 ANOVA that compared the freezing levels of the first three minutes of the SEFL trial prior to the shock in context B found three-way interactions between age, interval, and stress induction severity and between age, sex, and stress induction severity. This was followed by a 3-way ANOVA that looked at age, interval, and stress induction severity with sex collapsed.

| Generalized Freezing in Context B |  |  |  |
| --- | --- | --- | --- |
| Source | F | Sig. Value | Power |
| Age | 53.329 | < 0.001 | 1.000 |
| Sex | 20.676 | < 0.001 | 0.995 |
| Interval | .277 | 0.599 | 0.082 |
| Stress Induction Severity | 54.154 | < 0.001 | 1.000 |
| Age * Sex | 43.588 | < 0.001 | 1.000 |
| Age * Interval | 47.339 | < 0.001 | 1.000 |
| Age * Stress Induction Severity | 14.520 | < 0.001 | 1.000 |
| Sex * Interval | 11.040 | 0.001 | 0.912 |
| Sex * Stress Induction Severity | 8.487 | < 0.001 | 0.993 |
| Interval * Stress Induction Severity | .975 | 0.405 | 0.265 |
| Age * Sex * Interval | 1.430 | 0.233 | 0.222 |
| Age * Sex * Stress Induction Severity | 14.644 | < 0.001 | 1.000 |
| Age * Interval * Stress Induction Severity | 12.076 | < 0.001 | 1.000 |
| Sex * Interval * Stress Induction Severity | 1.152 | 0.329 | 0.309 |
| Age * Sex * Interval * Stress Induction Severity | .345 | 0.793 | 0.117 |

A 3-way ANOVA with sex collapsed confirmed an interaction between age, interval, and stress induction severity (F (3, 292) = 8.233, p < 0.001) (Table 15). This was followed by 2×4 ANOVAs that investigated the effects of stress induction severity and interval in adults and juveniles separately on generalized freezing in context B. Both analyses in juveniles and adults revealed an interaction between interval and stress severity (adult: F (3, 148) = 3.535, p < 0.05) (Table 16) (juvenile: F (3, 144) = 7.399, p < 0.001) (Table 17).

**Table 15:** Statistical outcomes for the 2×2×3 ANOVA with sex collapsed that looked at the interaction between age, interval, and stress induction severity on generalized freezing in context B, confirming a three-way interaction between age, interval, and stress induction severity (F (3, 292) = 8.233, p < 0.001).

| Generalized Freezing in Context B |  |  |  |
| --- | --- | --- | --- |
| Source | F | Sig. Value | Power |
| Age | 38.872 | < 0.001 | 1.000 |
| Interval | 0.150 | 0.698 | 0.067 |
| Stress Induction Severity | 38.378 | < 0.001 | 1.000 |
| Age * Interval | 31.464 | < 0.001 | 1.000 |
| Age * Stress Induction Severity | 10.876 | < 0.001 | 0.999 |
| Interval * Stress Induction Severity | 0.506 | 0.679 | 0.153 |
| Age * Interval * Stress Induction Severity | 8.233 | < 0.001 | 0.992 |

**Table 16:** Statistical outcomes for generalized freezing in context B for adults. A 2×4 ANOVA found an interaction between interval and stress induction severity. This was followed by a one-way ANOVA for each interval period that found a main effect of stress induction at both interval periods. *Post hoc* tests demonstrated a graded effect based on stress severity for both intervals.

| Generalized Freezing in Context B, Adults |  |  |  |
| --- | --- | --- | --- |
| Source | F | Sig. Value | Power |
| Interval | 11.9191 | < 0.001 | 0.929 |
| Stress Induction Severity | 29.698 | < 0.001 | 1.000 |
| Interval * Stress Induction Severity | 3.535 | < 0.001 | 0.776 |

| Generalized Freezing in Context B, Adult @ Recent Interval |  |  |  |
| --- | --- | --- | --- |
| Source | F | Sig. Value | Power |
| Stress Induction Severity | 9.316 | < 0.001 | 0.996 |
| *Post hoc Comparison* |  |  |  |
| Comparison |  | Mean Difference | Sig Value |
| No Stress | Low Stress | 1.645 | 1.000 |
| No Stress | Moderate Stress | 8.868 | 0.388 |
| No Stress | High Stress | 24.216 | < 0.001 |
| Low Stress | Moderate Stress | 7.223 | 0.921 |
| Low Stress | High Stress | 22.571 | < 0.001 |
| Moderate Stress | High Stress | 15.348 | 0.030 |

**Table 16:** Statistical outcomes for generalized freezing in context B for adults.
| Generalized Freezing in Context B, Adult @ Remote Interval |  |  |  |
| --- | --- | --- | --- |
| Source | F | Sig. Value | Power |
| Stress Induction Severity | 20.278 | < 0.001 | 1.000 |
| *Post hoc Comparison* |  |  |  |
| Comparison |  | Mean Difference | Sig Value |
| No Stress | Low Stress | 7.841 | 1.000 |
| No Stress | Moderate Stress | 24.016 | 0.003 |
| No Stress | High Stress | 49.644 | < 0.001 |
| Low Stress | Moderate Stress | 16.175 | 0.166 |
| Low Stress | High Stress | 41.803 | < 0.001 |
| Moderate Stress | High Stress | 25.629 | 0.005 |

**Table 17:**
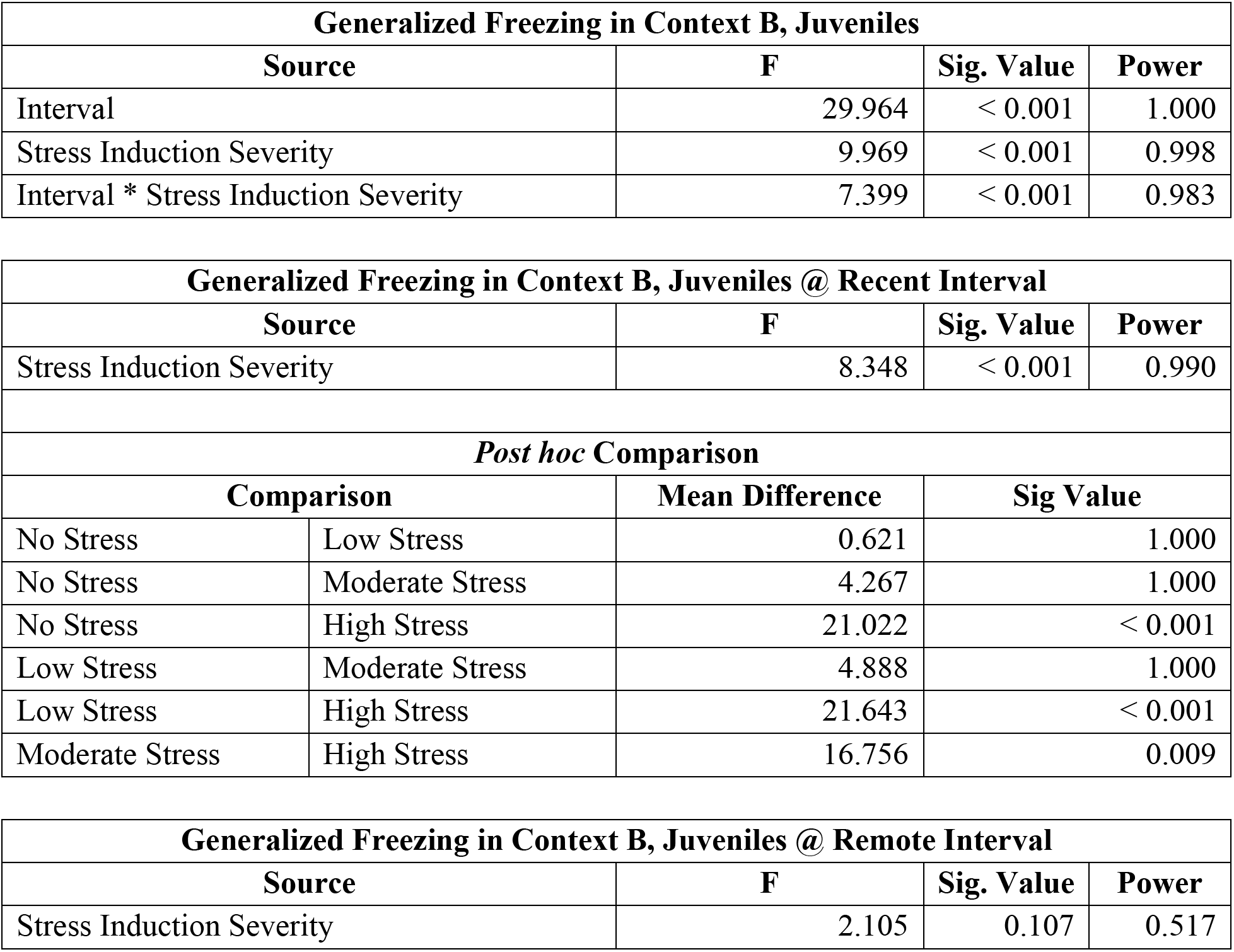
Statistical outcomes for generalized freezing in context B for juveniles. A 2×4 ANOVA found an interaction between interval and stress induction severity. This was followed by a one-way ANOVA for each interval period that found a main effect of stress induction at recent but not remote interval. *Post hoc* tests demonstrated a graded effect based on stress severity for the recent time point and no *post hoc* test was conducted for remote interval.

| Generalized Freezing in Context B, Juveniles |  |  |  |
| --- | --- | --- | --- |
| Source | F | Sig. Value | Power |
| Interval | 29.964 | < 0.001 | 1.000 |
| Stress Induction Severity | 9.969 | < 0.001 | 0.998 |
| Interval * Stress Induction Severity | 7.399 | < 0.001 | 0.983 |

| Generalized Freezing in Context B, Juveniles @ Recent Interval |  |  |  |
| --- | --- | --- | --- |
| Source | F | Sig. Value | Power |
| Stress Induction Severity | 8.348 | < 0.001 | 0.990 |
| *Post hoc Comparison* |  |  |  |
| Comparison |  | Mean Difference | Sig Value |
| No Stress | Low Stress | 0.621 | 1.000 |
| No Stress | Moderate Stress | 4.267 | 1.000 |
| No Stress | High Stress | 21.022 | < 0.001 |
| Low Stress | Moderate Stress | 4.888 | 1.000 |
| Low Stress | High Stress | 21.643 | < 0.001 |
| Moderate Stress | High Stress | 16.756 | 0.009 |

**Table 17:** Statistical outcomes for generalized freezing in context B for juveniles.
| Generalized Freezing in Context B, Juveniles @ Remote Interval |  |  |  |
| --- | --- | --- | --- |
| Source | F | Sig. Value | Power |
| Stress Induction Severity | 2.105 | 0.107 | 0.517 |

In adult rats at the recent interval, a follow-up ANOVA found a main effect of stress severity (F (3, 76) = 9.316, p < 0.001) (Figure 6A, Table 16) on pre-shock freezing in context B. Post hoc revealed this effect was driven by the high-stress group (p < 0.001), which froze significantly more than the no-stress group and low and moderate stress groups (low: p < 0.001, moderate: p < 0.05). Neither the low nor the moderate stress groups differed from the no-stress group (low: p = 1.000, moderate: p > 0.05). At the remote interval, a follow-up ANOVA in adult rats found a main effect of stress severity (F (3, 72) = 20.278, p < 0.001) (Figure 6B, Table 16). Post hoc comparisons showed that groups subjected to moderate (p < 0.01) and high (p < 0.001) stress levels exhibited significantly greater freezing behavior than the no-stress group; however, no significant difference was observed compared with the low-stress group (p > 0.05). This effect is partially graded to level of stress induction, as the high stress group exhibited greater freezing compared to the moderate (p < 0.01) and low (p < 0.001) stress groups; however, the moderate stress group did not significantly differ from the low stress group (p > 0.05).

**Figure 6:**
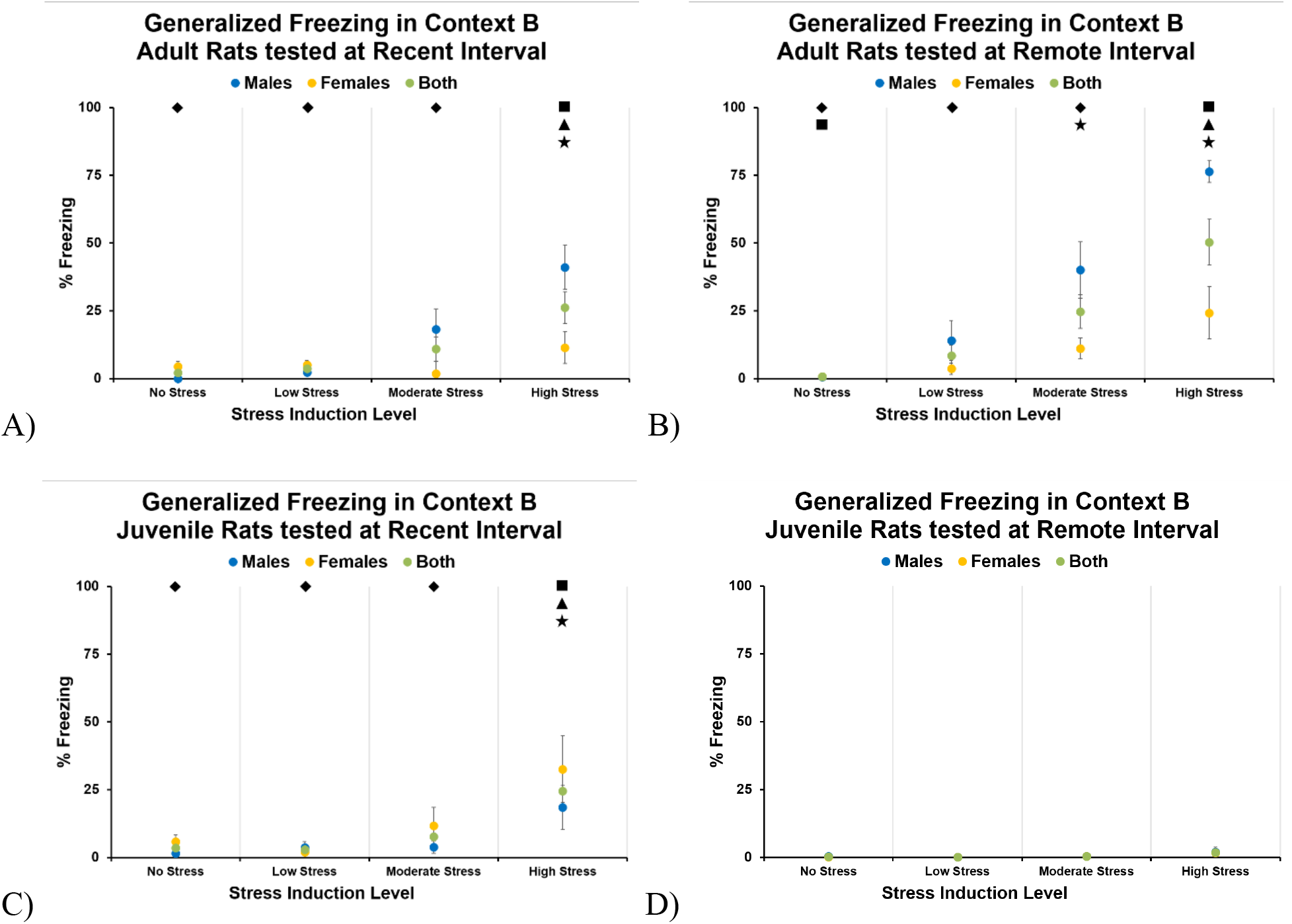
Pre-shock generalized freezing in context B. While no sex differences were found, all graphs depict male and female freezing levels along with the collapsed data. **A)** Adult rats when tested at recent interval found a main effect of stress, with high stress demonstrating higher generalized freezing than the other stress groups. **B)** Adult rats, when tested at remote interval, demonstrated shock-graded generalized freezing, with moderate and high stress differing from the no-stress group and each other. **C)** Juvenile rats, when tested at recent interval, showed an effect only for the high-stress group. **D)** Juvenile rats, when tested at remote interval, demonstrated no difference in generalized freezing based on stress induction severity.

In juvenile rats at the recent test interval, a subsequent ANOVA revealed a main effect of stress (F (3, 71) = 8.348, p < 0.001) (Figure 6C; Table 17). This effect resulted from the high-stress group, which exhibited greater freezing than the no-stress (p < 0.001), low-stress (p < 0.001), and moderate-stress (p < 0.01) groups. At the remote interval, juvenile rats, a follow-up ANOVA failed to reveal a significant effect of stress (F (1, 73) = 2.105, p > 0.05) (Figure 6D, Table 17)

### SEFL Test in Context B

To evaluate stress-enhanced fear learning, we assessed contextual fear retrieval in context B and 3 and 63 days after juvenile or adult stress induction in context A. A full-factorial 2×2×2×4 ANOVA found a four-way interaction among age, sex, interval, and stress-induction severity (F (3, 276) = 2.638, p = 0.05) (Table 18). This was followed by separate 2×4 ANOVAs to further evaluate how stress severity affected persistence in juvenile and adult male and female rats.

**Table 18:** Full factorial statistical outcomes for the SEFL test in context B. This 2×2×2×4 ANOVA found a four-way interaction between age, sex, interval, and stress induction severity. This was followed by 2×4 ANOVAs investigating interval and stress induction severity for adult males, adult females, juvenile males, and juvenile females.

| SEFL Test |  |  |  |
| --- | --- | --- | --- |
| Source | F | Sig. Value | Power |
| Age | 21.318 | < 0.001 | 0.996 |
| Sex | 3.442 | 0.065 | 0.456 |
| Interval | 6.730 | 0.010 | 0.734 |
| Stress Induction Severity | 86.004 | < 0.001 | 1.000 |
| Age * Sex | 8.490 | 0.004 | 0.827 |
| Age * Interval | 2.284 | 0.132 | 0.325 |
| Age * Stress Induction Severity | 1.458 | 0.226 | 0.384 |
| Sex * Interval | 0.632 | 0.427 | 0.124 |
| Sex * Stress Induction Severity | 0.426 | 0.734 | 0.135 |
| Interval * Stress Induction Severity | 2.813 | 0.040 | 0.673 |
| Age * Sex * Interval | 0.578 | 0.448 | 0.118 |
| Age * Sex * Stress Induction Severity | 1.122 | 0.341 | 0.301 |
| Age * Interval * Stress Induction Severity | 0.855 | 0.465 | 0.235 |
| Sex * Interval * Stress Induction Severity | 1.581 | 0.194 | 0.414 |
| Age * Sex * Interval * Stress Induction Severity | 2.638 | 0.050 | 0.642 |

Stress induction in adult male rats was partially graded and persisted (Figure 7A). A 2×4 ANOVA supported this, finding that there was a main effect of stress severity (F (3, 72) = 33.564, p < 0.001) (Table 19), but not of interval (F (1, 72) = 0.059, p > 0.05) nor an interaction between these variables (F (3, 72) = 0.369, p > 0.05). *Post hoc* comparisons confirmed this: the high (p < 0.001), moderate (p < 0.001), and low (p < 0.001) stress groups showed significantly greater freezing than the no-stress group. This enhancement showed modest evidence of stress severity-related grading, as the high-stress group was significantly greater than the low-stress group (p = 0.001) but did not significantly differ from the moderate-stress group (p > 0.05), nor did the low- and moderate-stress groups differ (p > 0.05).

**Figure 7:**
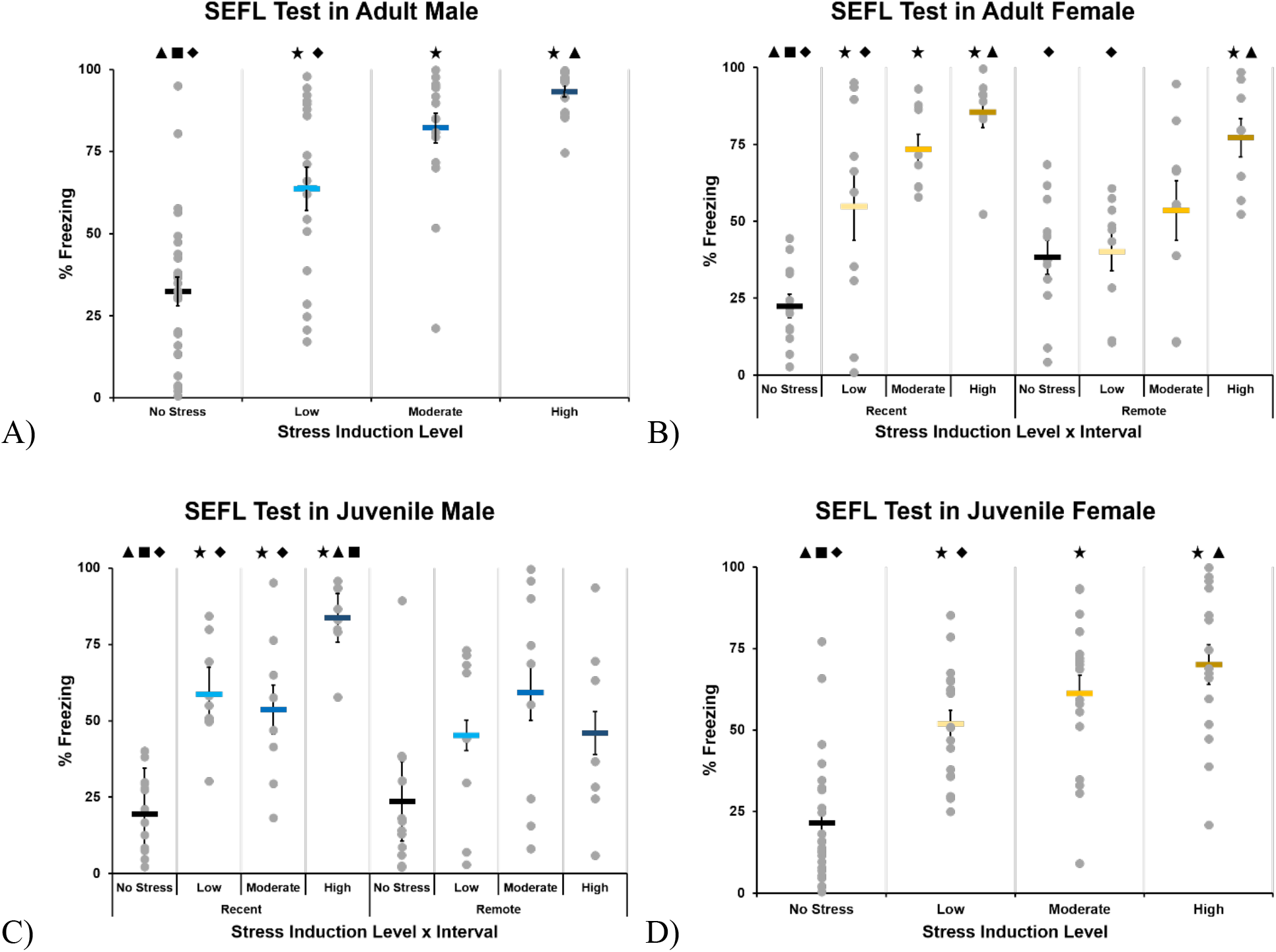
SEFL test following single-shock SEFL trial in context B. **A)** Adult males showed a graded SEFL that was influenced by stress induction severity. **B)** Adult females showed a graded enhancement when tested at recent interval, while at remote interval only high stress induction demonstrated enhanced fear learning compared to controls. **C)** Juvenile males demonstrated SEFL at recent interval, but not remote interval. This effect at recent interval is slightly graded, with low and moderate stress not differing from each other but significantly lower than the high-stress group. **D)** Juvenile females demonstrated enhancement of fear learning that was impacted only by stress severity, as all stress groups had higher freezing than the controls but not each other, meaning this effect is not graded.

**Table 19:**
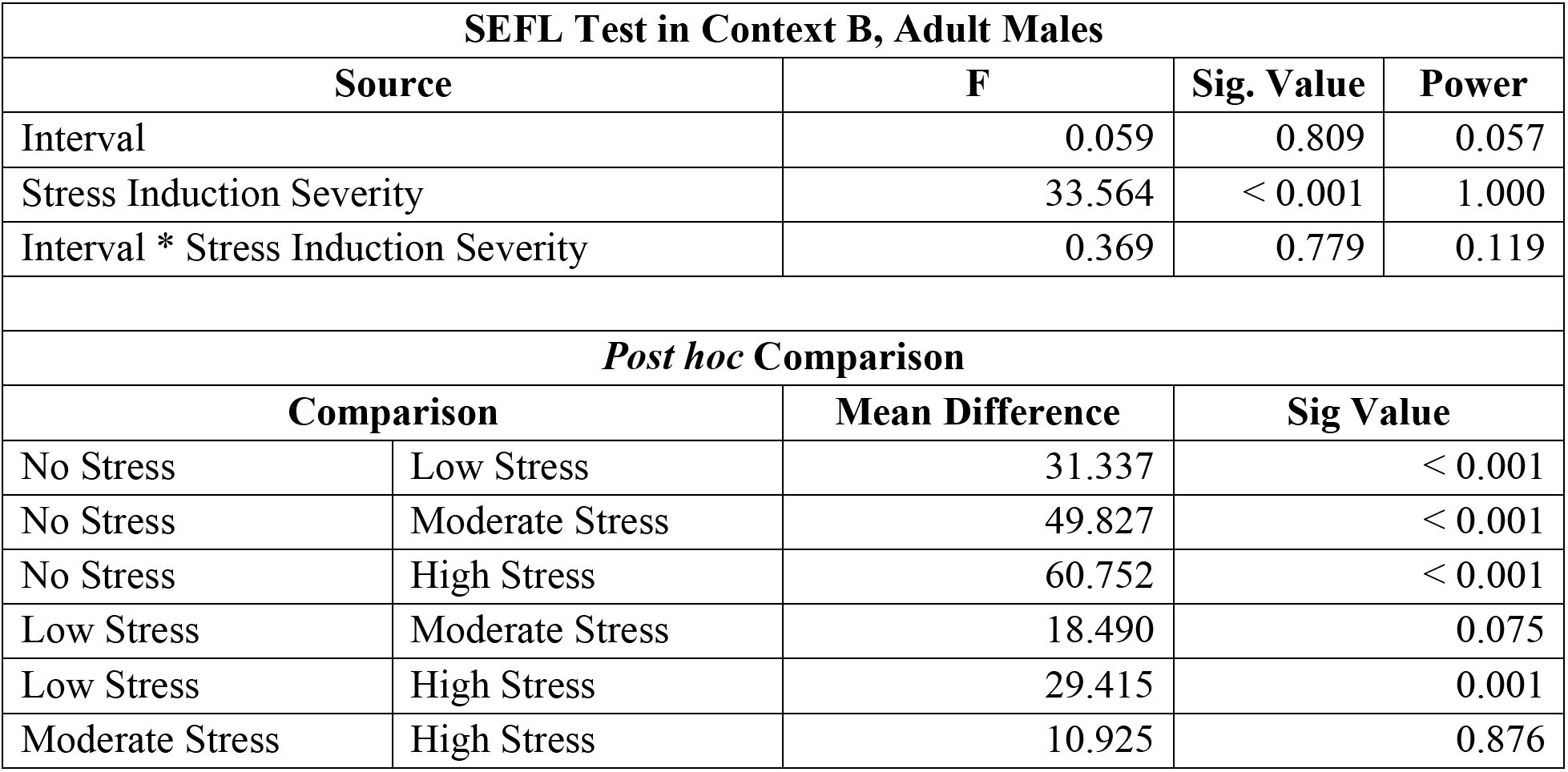
Statistical outcomes for the SEFL test in context B, adult males. Only a main effect of stress induction severity was found. A *post hoc* comparison between stress groups confirmed that stress induction enhanced context fear learning across all stress induction groups as the high, moderate, and low stress groups exhibited significantly greater freezing than the control group.

Stress induction in adult females was affected by both stress severity and its post-interval duration (Figure 7B), as supported by a 2×4 ANOVA that found a significant interaction (F (3, 68) = 2.891, p < 0.05). This was followed by one-way ANOVAs that found a main effect of stress severity in the recent (F (3, 34) = 16.512, p < 0.001) and remote (F (3, 34) = 6.048, p < 0.01) periods (Table 20). A *post hoc* comparison between stress groups at the recent test period confirmed that stress induction enhanced context fear learning across all stress severity groups, as the high (p < 0.001), moderate (p < 0.001), and low (p < 0.01) stress groups exhibited significantly greater freezing than the non-stressed group. We also see that this enhancement is graded based on the degree of stress induction, as the high-stress group was significantly greater than the low-stress group (p < 0.05), but the moderate-stress group was not significantly different from either the high (p = 1.000) or low (p > 0.05) stress group. At the remote test period, *post hoc* comparisons between stress groups confirmed that stress induction enhanced context fear learning only in the high-stress group (p < 0.01), which exhibited significantly greater freezing than the non-stressed group.

**Table 20:** Statistical outcomes for the SEFL test in context B, adult females. A 2×4 ANOVA found an interaction between interval and stress induction severity which was followed by one-way ANOVAs at both interval periods that found a main effect of stress induction severity at the recent and remote interval time point. A *post hoc* comparison for the recent time point confirmed that stress induction enhanced context fear learning across all stress induction severity groups in a graded effect. While a *post hoc* comparison for the remote interval found that only the high stress group displayed significantly higher freezing than the control group.

| SEFL Test in Context B, Adult Females |  |  |  |
| --- | --- | --- | --- |
| Source | F | Sig. Value | Power |
| Interval | 1.792 | 0.185 | 0.262 |
| Stress Induction Severity | 19.722 | < 0.001 | 1.000 |
| Interval * Stress Induction Severity | 2.891 | 0.042 | 0.665 |

| SEFL Test in Context B, Adult Females @ Recent Interval |  |  |  |
| --- | --- | --- | --- |
| Source | F | Sig. Value | Power |
| Stress Induction Severity | 16.512 | < 0.001 | 1.000 |
| *Post hoc* Comparison |  |  |  |
| Comparison |  | Mean Difference | Sig Value |
| No Stress | Low Stress | 32.355 | 0.008 |
| No Stress | Moderate Stress | 61.009 | < 0.001 |
| No Stress | High Stress | 63.091 | < 0.001 |
| Low Stress | Moderate Stress | 18.654 | 0.457 |
| Low Stress | High Stress | 30.736 | 0.029 |
| Moderate Stress | High Stress | 12.082 | 1.000 |

**Table 20:** Statistical outcomes for the SEFL test in context B, adult females.
| SEFL Test in Context B, Adult Females @ Remote Interval |  |  |  |
| --- | --- | --- | --- |
| Source | F | Sig. Value | Power |
| Stress Induction Severity | 6.048 | 0.002 | 0.936 |
| *Post hoc* Comparison |  |  |  |
| Comparison |  | Mean Difference | Sig Value |
| No Stress | Low Stress | 1.829 | 1.000 |
| No Stress | Moderate Stress | 15.183 | 0.731 |
| No Stress | High Stress | 38.908 | 0.002 |
| Low Stress | Moderate Stress | 13.354 | 1.000 |
| Low Stress | High Stress | 37.078 | 0.008 |
| Moderate Stress | High Stress | 23.725 | 0.186 |

Stress induction in juvenile males was robust at the recent interval but was not confirmed at the remote interval (Figure 7C). A 2×4 ANOVA showed an interaction between interval and stress severity (F (3, 69) = 3.143, p < 0.05). This was followed by one-way ANOVAs that found a main effect of stress induction at the recent (F (3, 36) = 28.709, p < 0.001) but not the remote (F (3, 33) = 2.824, p > 0.05) post-stress period (Table 21). In the recently tested groups, a *post hoc* comparison between stress groups at recent interval confirmed that stress induction enhanced context fear learning across all stress induction groups, as the high (p < 0.001), moderate (p < 0.001), and low (p < 0.001) stress groups exhibited significantly greater freezing than the non-stressed group. This enhancement was partially graded based on the degree of stress induction. The high-stress group was significantly greater than the moderate (p < 0.01) and low (p < 0.05) stress groups, but the moderate-stress group did not differ significantly from the low-stress group (p = 1.000).

**Table 21:**
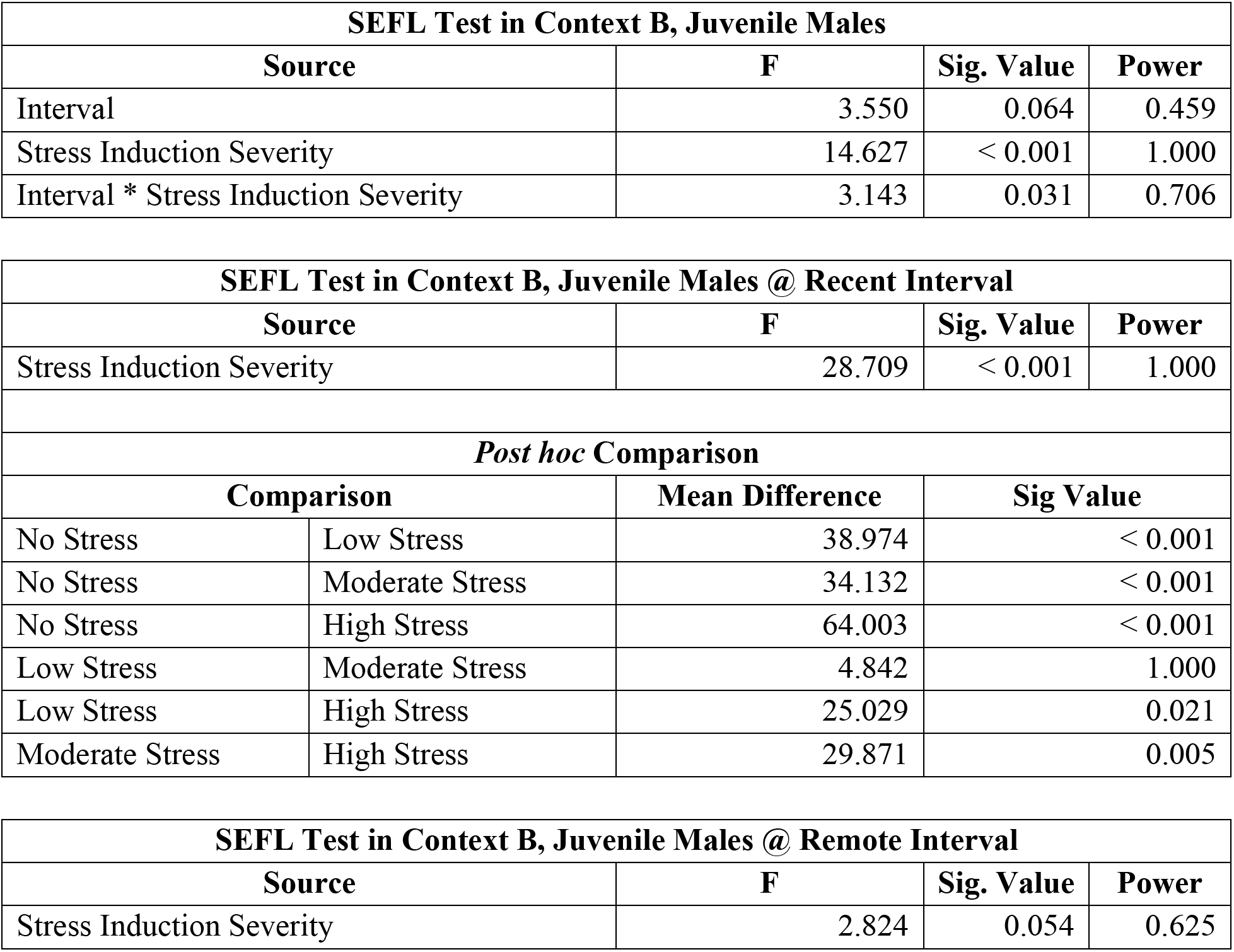
Statistical outcomes for the SEFL test in context B, juvenile males. A 2×4 ANOVA found an interaction between interval and stress induction severity. This was followed by one-way ANOVAs that found a main effect of stress induction at recent but not remote interval. A *post hoc* between stress groups at recent interval confirmed that stress induction enhanced context fear learning across all stress induction groups as high, moderate, and low stress groups exhibited significantly higher freezing than the no stressed group. No *post hoc* comparison was conducted on stress groups at remote interval.

Stress-induction in juvenile females demonstrated a robust and persistent SEFL (Figure 7D) as supported by a 2×4 ANOVA, which identified main effects of interval (F (1, 67) = 4.720, p < 0.05) and stress severity (F (3, 67) = 23.771, p < 0.001), but no interaction these conditions (F (3, 67) = 1.022, p > 0.05) (Table 22). A *post hoc* comparison between stress groups confirmed that stress induction enhanced context fear learning across all stress induction groups, as the high (p < 0.001), moderate (p < 0.001), and low (p < 0.001) stress groups exhibited significantly greater freezing than the non-stressed group, while no significant differences were found between low, moderate, or high stress conditions.

**Table 22:**
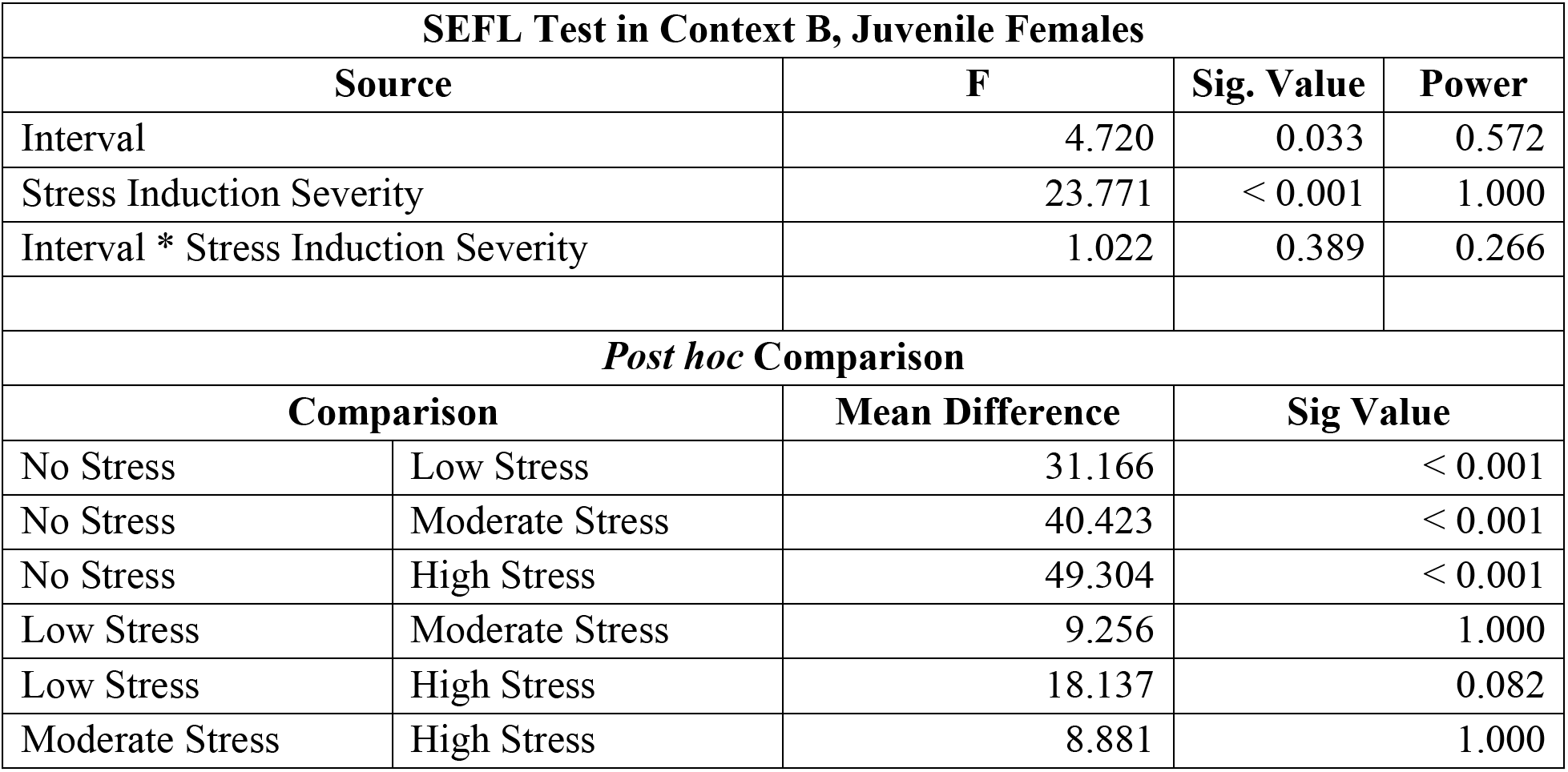
Statistical outcomes for the SEFL test in context B, juvenile females. A 2×4 ANOVA found only a main effect of stress induction severity was found. A *post hoc* comparison between stress groups confirmed that stress induction enhanced context fear learning across all stress induction groups as the high, moderate, and low stress groups exhibited significantly greater freezing than the control group. This effect was not graded as none of the stress groups were found to be significantly different from each other.

### % Change SEFL Test in Context B

To compare SEFL expression between sex and age groups and account for differences in freezing levels related to these variables in control groups, we calculated percent change for each group relative to its respective controls. Stress induction in adult males resulted in persistent SEFL, while adult females showed decreased SEFL (Figure 8A). Conversely, stress induction during the juvenile period reversed the effect, with juvenile females showing persistence and juvenile males showing decreased SEFL (Figure 8B). This was supported by a significant interaction between age, sex, and interval (F (1, 179) = 27.088, p < 0.001) (Table 23). This was followed up by an ANOVA investigating sex and interval interactions for juvenile and adult stress groups separately, with stress severity collapsed. Stress during adulthood resulted in an interaction between sex and test interval (F (1, 100) = 21.193, p < 0.001) (Table 24). This cross-over interaction indicates an adult sex difference in the persistence of stress enhancement, with females showing a significant reduction in freezing at the remote than the recent test interval, whereas in males freezing did not differ across this same interval. Conversely, stress induction during the juvenile period resulted in an interaction between sex and interval (F (1, 95) = 6.585, p < 0.05) (Table 25); males exhibited a significant reduction at the remote test period compared with the recent period, while in females freezing did significantly differ between these intervals.

**Figure 8:**
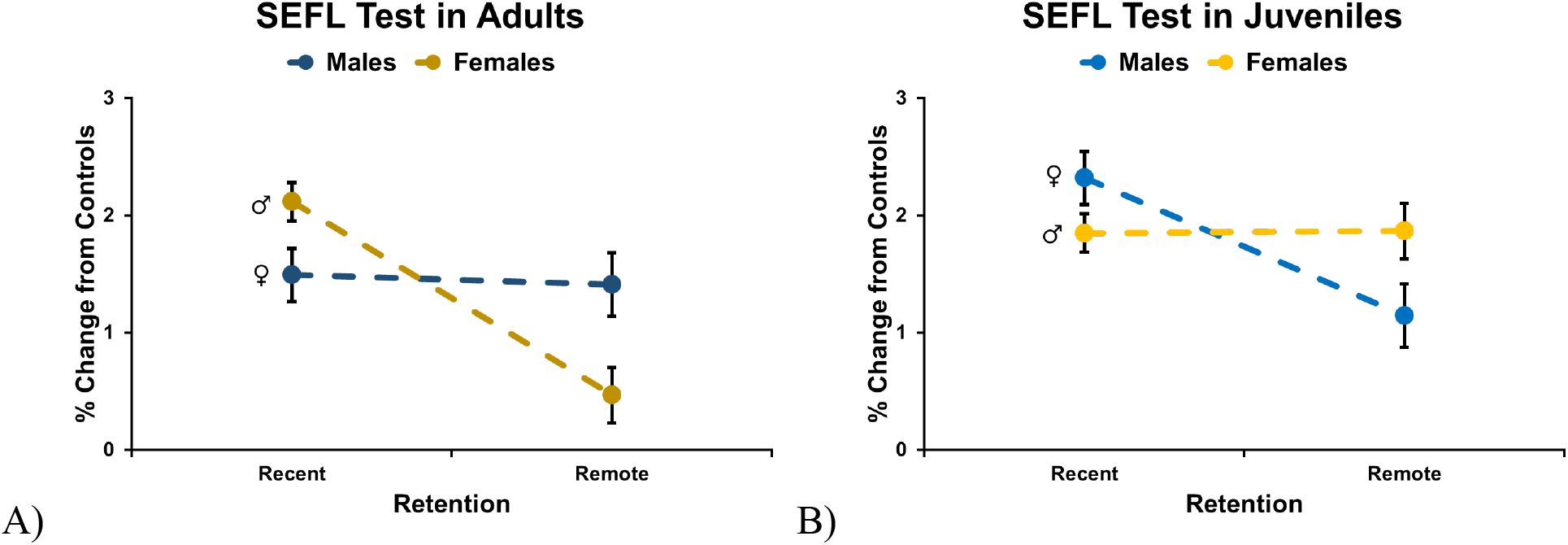
Percent change from controls for SEFL testing 1 day following single-shock SEFL trial in Context B. An interaction between age, sex, and interval was found, allowing for the collapse of stress induction severity. A two-way ANOVA for each developmental age found an interaction between sex and test interval, with a crossover effect. A) Adult male rats demonstrated a persistence of fear enhancement across interval periods, while adult female rats showed a decrease in fear enhancement over time. B) Juvenile male rats demonstrated a decrease in fear enhancement between interval periods, while juvenile females demonstrated a robust persistence of fear enhancement over time.

**Table 23:** Full factorial statistical outcomes for the % change SEFL test in context B. To compare SEFL expression between sex and age groups and to account for difference in freezing levels in control groups, the percent change was calculated for each group from its respective controls. A 2×2×2×3 ANOVA found an interaction between age, sex, and interval. This was followed up with a 2×2 ANOVAs investigating sex and interval interactions at juvenile and adult developmental ages separately with stress levels collapsed.

| % Change SEFL Test in Context B |  |  |  |
| --- | --- | --- | --- |
| Source | F | Sig. Value | Power |
| Age | 9.182 | 0.003 | 0.854 |
| Sex | 0.004 | 0.952 | 0.050 |
| Interval | 32.015 | < 0.001 | 1.000 |
| Stress Induction Severity | 14.941 | < 0.001 | 0.999 |
| Age * Sex | 1.157 | 0.283 | 0.188 |
| Age * Interval | 1.178 | 0.279 | 0.190 |
| Age * Stress Induction Severity | 0.481 | 0.619 | 0.128 |
| Sex * Interval | 0.475 | 0.491 | 0.105 |
| Sex * Stress Induction Severity | 0.286 | 0.751 | 0.095 |
| Interval * Stress Induction Severity | 1.373 | 0.256 | 0.293 |
| Age * Sex * Interval | 27.088 | < 0.001 | 0.999 |
| Age * Sex * Stress Induction Severity | 0.084 | 0.920 | 0.063 |
| Age * Interval * Stress Induction Severity | 0.739 | 0.479 | 0.174 |
| Sex * Interval * Stress Induction Severity | 2.187 | 0.115 | 0.443 |
| Age * Sex * Interval * Stress Induction Severity | 2.539 | 0.082 | 0.503 |

**Table 24:** Statistical outcomes for the % change SEFL test in context B, adults. A 2×2 ANOVA found an interaction between sex and interval, a main effect of interval, but not sex indicating a cross-over interaction.

| % Change SEFL Test in Context B, Adults |  |  |  |
| --- | --- | --- | --- |
| Source | F | Sig. Value | Power |
| Sex | 0.866 | 0.354 | 0.152 |
| Interval | 25.888 | < 0.001 | 0.999 |
| Sex * Interval | 21.193 | < 0.001 | 0.995 |

**Table 25:**
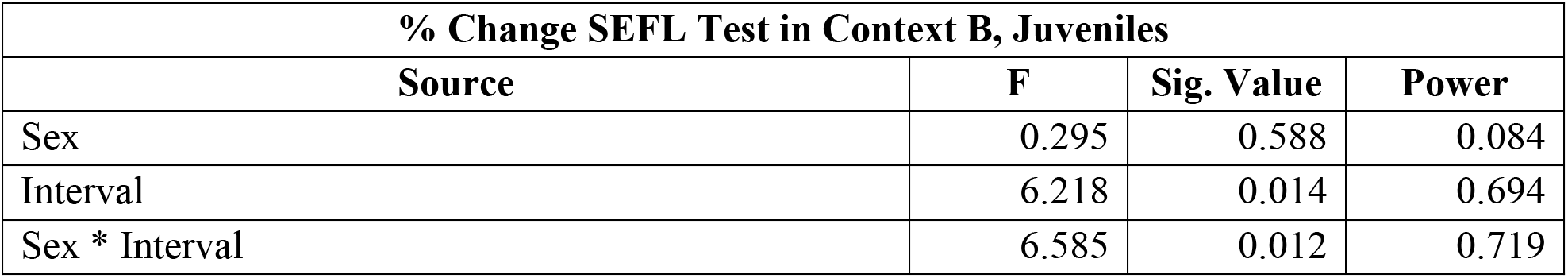
Statistical outcomes for the % change SEFL test in context B, juveniles. A 2×2 ANOVA found an interaction between sex and interval, a main effect of interval, but not sex indicating a cross-over interaction.

| % Change SEFL Test in Context B, Juveniles |  |  |  |
| --- | --- | --- | --- |
| Source | F | Sig. Value | Power |
| Sex | 0.295 | 0.588 | 0.084 |
| Interval | 6.218 | 0.014 | 0.694 |
| Sex * Interval | 6.585 | 0.012 | 0.719 |

## Discussion

The present study examined how the severity of an acute stress experience during the juvenile and adult periods influences the establishment, persistence, and magnitude of stress-enhanced fear learning (SEFL) in male and female rats. By systematically varying the number of footshocks during stress induction and assessing the persistence of stress-induced memory and the subsequent enhancement of recent and remote fear learning, we made several important findings with potential implications for posttraumatic stress disorder. First, although all groups showed stress-enhanced learning shortly after stress induction, maintenance of SEFL depended on sex and the developmental timing of stress induction. In females, only the most severe stress induction (15 footshocks) during adulthood led to a remote enhancement of fear learning. Whereas in males, stress induction during the juvenile period led to a bimodal pattern of fear learning that did not reach significance. This pattern was consistent with the relative magnitude measures of SEFL, in which SEFL strength declined over time when stress induction occurred in juvenile males (but not in juvenile females) and in adult females (but not in adult males). Second, stress-induced contextual fear memory and SEFL generally increased as a function of stress severity, indicating that traumatic stress produces graded rather than all-or-none changes in threat learning. Third, adult animals exhibited highly persistent contextual fear memories that remained stable across a two-month retention interval, whereas juvenile animals showed developmental constraints on the long-term persistence of stress memories. Finally, changes in contextual fear memory did not always parallel changes in SEFL, suggesting that memory for the traumatic event and sensitized fear learning are dissociable.

An important consideration in interpreting the current findings was whether group differences in SEFL could be explained by variations in initial shock sensitivity, associative learning during stress induction, or subsequent differences in fear memory retention and generalization. Reactivity to footshock did not differ across age or sex groups, indicating that the observed behavioral effects are unlikely to reflect developmental or sex-dependent differences in nociceptive reactivity. Furthermore, under both moderate- and high-stress conditions, associative fear learning progressed similarly across groups, as evidenced by an increase in post-shock freezing with repeated shock exposure. Neither age nor sex significantly influenced this measure. Assessments of contextual fear memory further support this interpretation while accounting for developmental differences in long-term stress memory retention. Adult males and females exhibited robust, graded contextual fear retention that remained stable across the 60-day retention interval. In contrast, juvenile males and females showed robust recent contextual fear but reduced retention at remote time points, consistent with our previous findings (Odynocki et al., 2023). Patterns of fear generalization also differed across groups. Adult males exhibited substantial freezing in a novel context at both recent and remote intervals despite moderate extinction, whereas juvenile females showed generalized freezing only at the recent test interval. Although the relationship between fear generalization and SEFL at the remote interval was apparent in adult males and females, it was absent in juvenile females. Importantly, previous work has demonstrated that extinction or generalization prior to SEFL trial in context B is not a significant determinant of SEFL expression (Long & Fanselow, 2012). Therefore, differences in associative fear retention or generalization are unlikely to account for the temporal decline in SEFL observed in juvenile males and adult females. Collectively, these results suggest that developmental and sex-dependent differences in SEFL are more likely attributable to nonassociative stress memory processes than to variations in associative fear memory or in the unconditional response to footshock.

The central finding of this study concerns the persistence of SEFL. Consistent with previous reports, prior stress exposure enhanced subsequent learning to a single-shock threat. Importantly, this enhancement was graded by stress severity, which may indicate that nonassociative sensitization processes scale with the severity of the initial stress experience or the number of stimulus presentations. This observation extends prior SEFL findings (Rau et al., 2005) by demonstrating that relatively mild stressors (a single footshock) can produce enhancements, whereas increasingly severe stressors produce progressively larger effects. Such graded relationships are important because traumatic experiences in natural situations vary substantially in severity.

The persistence of SEFL differed substantially across age and sex groups. Adult males exhibited perhaps the most stable expression, showing robust enhancements in fear learning at both recent and remote intervals, as previously reported (Rau & Fanselow, 2009). These findings indicate that the sensitized state produced by acute stress in adult rats can persist for at least three months without decline. Adult females exhibited a distinct pattern of SEFL persistence. While robust SEFL was observed shortly after stress induction as previously described (Poulos et al., 2015), only the highest stress level produced a significant enhancement at the remote interval. Moreover, analysis of relative SEFL magnitude indicated a pronounced decline across time. Interestingly, females displayed stronger relative SEFL than males at recent time points, yet a weaker persistence at remote time points. Both results are novel, with the former result seemingly in conflict with prior work (Hassien et al., 2020; Gonzalez et al., 2021; Poulos et al., 2014); however, none of the studies considered non-shock control levels of context B fear learning to establish a magnitude measure of SEFL under conditions that correct for sex- and developmental differences in contextual fear learning (Colon et al., 2018). The latter finding, whether using raw or relative SEFL measures, reveals a considerable decline in SEFL persistence with lower-stress induction procedures in females. This pattern suggests that the mechanisms governing the induction of stress sensitization may differ from those responsible for its maintenance. Female animals may initially develop a stronger sensitized state but exhibit a moderation over time. Indeed, fluctuations in ovarian hormone signaling and sex-specific stress neurobiology may contribute to this time-dependent SEFL profile.

The juvenile groups revealed a potentially more complex developmental pattern. Stress in juvenile males showed a robust recent SEFL but failed to maintain this enhancement when tested at a remote interval as adults. In contrast, juvenile females exhibited persistent SEFL across both intervals. Particularly striking, when examining SEFL magnitude, the crossover interaction observed between sex and interval in juveniles suggests that the developmental timing of stress and subsequent development across this long interval alters how sex or sex hormones influence stress-induced plasticity. Recently, Quinn and colleagues (2024) found that stress induction at an earlier period in 17-day-old rat pups, using 4 but not 15 footshocks, produced SEFL after 74 days in females but not in males. These findings are particularly intriguing because they imply that the mechanism supporting long-term sensitization is present in prepubertal females but not in prepubertal males, or that males may be particularly affected by pubertal gonadal hormones. Such developmental divergence may reflect differences in the maturation of stress-responsive neural circuits, hormonal influences on plasticity, or sex-specific trajectory of amygdala-prefrontal or hippocampal development.

Perhaps a more theoretically important observation is the partial dissociation between contextual fear memory and SEFL. In the current study, the persistence of context fear did not predict the persistence of sensitized fear learning, an observation previously reported (Poulos et al., 2014; Quinn et al., 2014). For example, contextual fear memories remained detectable in some groups despite reductions in SEFL, whereas other groups showed persistent SEFL despite developmentally mediated deficits in retrieving context fear memories. These findings support the growing view that SEFL is not simply a byproduct of strong traumatic memory (Poulos et al., 2014; Rau et al., 2005). Instead, traumatic stress may engage at least two partially independent processes: one responsible for storing trauma memories and another that alters the future sensitivity of fear learning systems.

At a neural level, this distinction may reflect differential involvement of hippocampal, prefrontal cortical, and amygdalar circuitry. Context fear memories, clearly established in adults, depend on hippocampal representations of the contextual environment, whereas SEFL has been linked more closely to stress-induced plasticity within amygdala-centered learning networks. Indeed, intraventricular injections of the NMDA antagonist AP5 disrupt hippocampal-dependent context fear memories, but fail to disrupt SEFL (Rau et al., 2005). Conversely, amygdala-targeted infusions of the GABA-A receptor agonist muscimol disrupt both contextual fear memory and SEFL (Perusini et al., 2016). Acute stress or trauma may therefore produce enduring modifications or plasticity in the amygdala, via corticosterone-mediated neuromodulation or stress-responsive molecular pathways that persist independently of the original contextual memory. Such mechanisms could explain why the magnitude and persistence of sensitization do not always mirror the retention of context fear, suggesting potential nonassociative processes.

Despite the comprehensive experimental design, some limitations should be recognized to guide future studies. First, sex differences in adult stress responses during SEFL testing were not assessed; therefore, the contributions of stress hormones at these later times remain poorly understood. Second, the study focused on behavioral measures, limiting the ability to identify the neural mechanisms underlying developmental and sex-dependent differences. Future work that times stress induction and SEFL tests during specific phases of the estrus cycle may have important implications for SEFL persistence. Finally, work incorporating immediate-early gene mapping, circuit manipulations, and CA^+2^ imaging approaches could determine how the developmental status of the fear network at the time of stress induction, and how stress induction alters the function of this network, will be important.

The present findings provide a wide-ranging characterization of how stress severity, developmental stage, sex, and the passage of time interact to shape both stress/traumatic memory and SEFL. These results demonstrate that the consequences of acute trauma are graded, enduring, and highly dependent on key biological variables such as sex and age. Adults exhibit robust sensitization, whereas developmental stage and sex strongly influence the long-term trajectory of stress-induced plasticity. The dissociation between stress memory retention and subsequent sensitized learning suggests that trauma produces enduring changes in the learning system that extend beyond memory of the original event. Understanding the mechanisms that support these long-lasting alterations may provide important insight into the developmental origins of vulnerability and resilience following traumatic stress.

